# Branched chain fatty acid-rich diet promotes lipid droplet enlargement and impacts organismal health in *Caenorhabditis elegans*

**DOI:** 10.1101/2025.01.15.633283

**Authors:** Kam Ue Roy Li, Wing Ka Lo, Meigui Yang, Chenyin Wang, Lau Chun Yin, Jetty Chung-Yung Lee, Chaogu Zheng

**Affiliations:** School of Biological Sciences, The University of Hong Kong, Hong Kong SAR, China

## Abstract

Considerable amounts of branched chain fatty acids (BCFAs) are present in the human diet from beef and dairy products. BCFAs can also be produced by the human gut microbiota and synthesized from branched chain amino acids. However, the physiological impact of a BCFA-rich diet on lipid metabolism and organismal health is unclear. In this study, by screening a collection of dietary bacteria, we find that the BCFA-rich *Microbacterium* diet causes the formation of supersized LDs and delays development, reduces brood size, and shortens lifespan of *C. elegans*. The high-BCFA diet downregulates *argk-1/*creatine kinase to inhibit the AMPK pathway and β-oxidation and upregulates *fat-7*/desaturase to promote the accumulation of PUFA, which enhances lipogenesis and LD expansion. We also isolate a gain-of-function mutation in *scav-4*/CD36, which enhances BCFA absorption and exacerbates BCFA-induced LD enlargement, demonstrating that host genetic variation in a fatty acid transporter could influence the susceptibility to a high-BCFA diet.

## INTRODUCTION

Lipids play an essential role in energy homeostasis, cell membrane construction, and signaling transduction, thus contributing to the development, physiology, and function of an organism. Dysregulation of lipid metabolism is associated with a range of health problems, such as obesity, cardiovascular diseases, and diabetes. Excessive lipid accumulation can cause non-alcoholic fatty liver disease and atherosclerosis, while impaired lipogenesis may lead to lipodystrophy and cachexia.^1^ Although dietary and genetic factors contribute significantly to the individual variation in lipid metabolism and susceptibility to lipid disorders,^2^ it remains challenging to understand how the lipid composition of the diet interacts with the genetic makeup of the host. Moreover, recent studies have also identified an emerging role of the gut microbiota in modulating metabolic phenotypes,^3^ suggesting a potential three-way interaction among the host, diet, and gut microbiome in regulating lipid metabolism and energy balance.

For example, branched chain fatty acids (BCFAs) are saturated fatty acids with one or more methyl branches on the carbon chain. In monomethyl BCFAs, the methyl group is usually attached to the penultimate (*iso*) or the antepenultimate (*anteiso*) carbon atom.^4^ BCFAs are naturally synthesized in many gram-positive bacteria, such as *Bacillus, Streptomycetes,* and *Staphylococcus*.^5^ Human diet contains BCFAs (∼400 mg/day in American adults) from dairy products and beef because BCFAs are produced by bacteria in the rumen of cattles.^6^ Interestingly, the human gut microbiota also produce BCFAs, and the production is influenced by food intake and diet-mediated changes in the gut microbiome.^7^ In addition, BCFAs can be *de novo* synthesized from the catabolic products of branched chain amino acids (BCAAs).^8^ Despite their presence in the human body, the biological function of BCFAs is not entirely understood. Previous studies found that BCFAs found in the neonatal gut can reduce the incidence of necrotizing enterocolitis,^9^ and exert cytotoxicity on breast cancer cells^10^ and multiple other human cancer cell lines,^11^ suggesting certain beneficial effects on human health. However, the physiological impact of a BCFA-rich diet on lipid metabolism and organismal health is unclear. Moreover, the genetic variants of the host that can influence the metabolic effects of dietary BCFAs are unknown.

In this study, we investigated how a high-BCFA diet affects lipid droplet size, energy balance, development, reproduction, and lifespan of the nematode *Caenorhabditis elegans*, which has recently emerged as an excellent model for understanding the mechanisms of host-diet-microbe interactions.^12^ *C. elegans* forms lipid droplets (LDs) in the intestine and hypodermis to store excessive fat in a process similar to the LD formation in mammalian adipose tissue. In *C. elegans*, the size of LDs is tightly controlled by multiple genetic and metabolic pathways.^13^ Mutations in genes encoding triglyceride synthesis complex,^14^ seipin,^15^ stearoyl-CoA desaturase,^16^ and atlastin GTPases^17^ reduce LD size, whereas mutations in genes involved in peroxisomal import of fatty acids, peroxisomal β-oxidation,^18,19^ and phosphatidylcholine synthesis^20^ increase LD size. Dietary fatty acids also regulate the formation of LDs. Monounsaturated fatty acids (MUFAs) upregulate the number but not the size of LDs,^21^ while polyunsaturated fatty acids (PUFAs) and microbial cyclopropane fatty acids (CFAs) promote seipin enrichment on LDs and the expansion of LD size.^15^ The standard laboratory food for *C. elegans* is *Escherichia coli* OP50, but anecdotal studies have shown that other dietary bacteria, such as *E. coli* HT115 and HB101,^22^ or pathogenic bacterium, such as *Stenotrophomonas maltophilia*, could alter LD size,^23^ suggesting potential microbial regulation of host metabolism.

*C. elegans* in the wild likely feed on a diverse range of bacterial species other than *E. coli*, and previous studies have isolated hundreds of these natural bacteria strains (belonging to many different phyla) from the natural habitat of the *C. elegans*.^24^ To understand whether these natural bacterial diets influence *C. elegans* lipid metabolism, we screened a collection of 429 such strains for increased LD size and identified several *Microbacterium* strains whose lipids are mostly BCFAs. We found that the abundant dietary BCFAs promote the formation of supersized LDs, delayed development, reduced brood size, and shortened lifespan of *C. elegans*. Transcriptomic analysis revealed that a BCFA-rich diet led to the downregulation of *argk-1*/creatine kinase and subsequent inhibition of AMPK pathway, as well as the accumulation of PUFAs through the upregulation of *fat-7*/desaturase. These signaling changes resulted in suppressed β-oxidation and enhanced lipogenesis and LD expansion. Moreover, through a genetic screen, we isolated a gain-of-function mutation in *scav-4* (a *C. elegans* homolog of CD36), which enhanced BCFA absorption and exacerbated the effects of a BCFA-rich diet on LD enlargement. Thus, this study not only elucidated the mechanisms by which dietary BCFAs promote lipid accumulation but also demonstrated that host genetic variation in a fatty acid transporter could influence susceptibility to a high-BCFA diet.

## RESULTS

### *Microbacterium* diet increased lipid droplet size in *C. elegans*

By feeding a collection of natural bacteria^24^ individually to *C. elegans* and observing the LD size in the intestine of adults, we identified a bacteria strain called JUb74, which would induce the formation of enlarged lipid droplets, especially in the posterior intestine (Figure 1A-C). The identity of these lipid droplets was confirmed by staining with C1-BODIPY-C12 in live animals (Figure 1B) and by the colocalization with the intestinal lipid droplet markers MDT-28::mCherry, DHS-3::GFP, and mRuby::DGAT-2 (Figure 1D and S1A). Most LDs in the animals fed with *E. coli* OP50 were smaller than 3 μm in diameter with a few reaching 3-4 μm, whereas the animals fed with JUb74 had many more 3-4 μm LDs and larger ones with the diameter ranging from 4 to 7 μm (Figure 1C).

**Figure 1.**
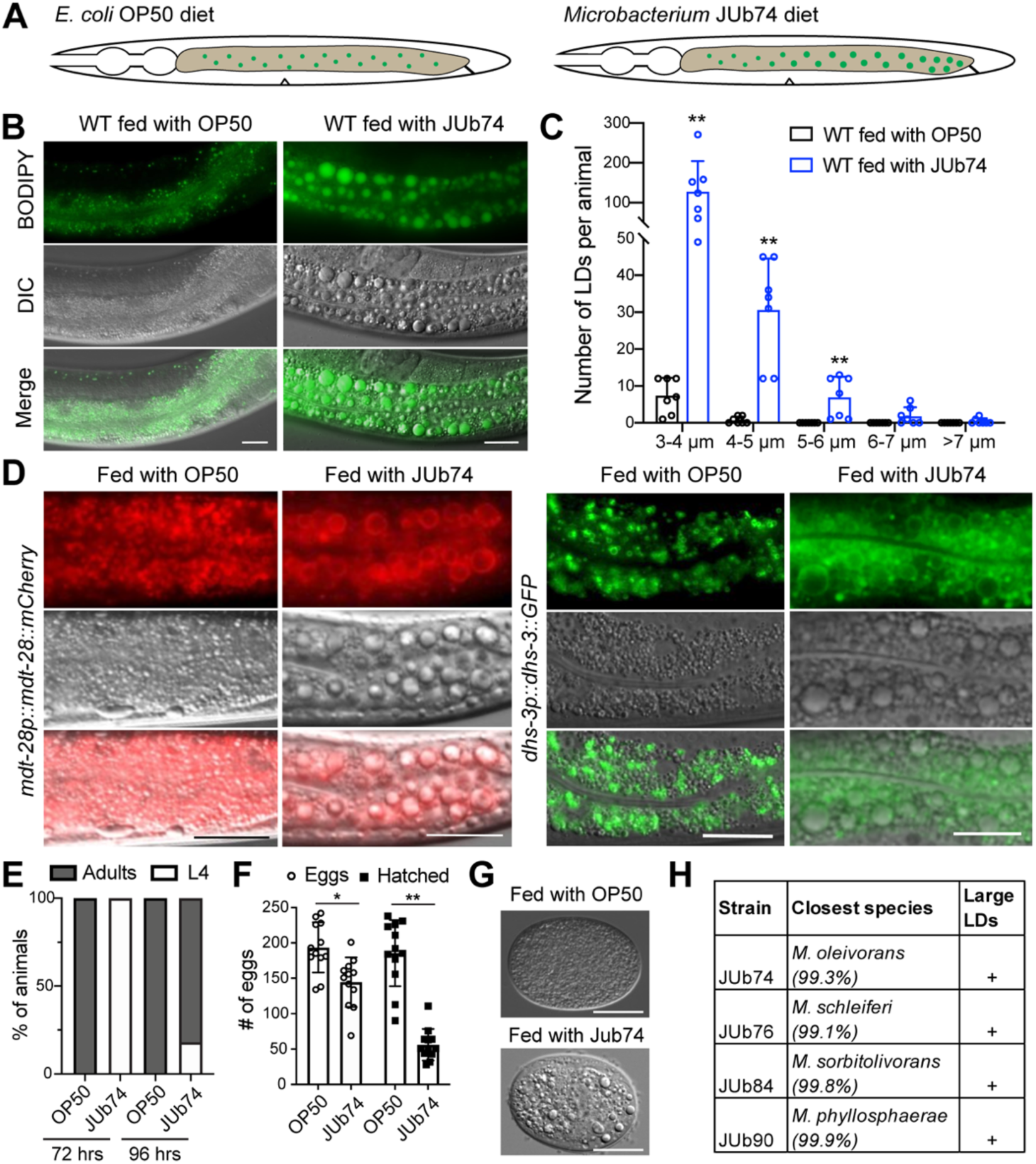
JUb74 diet causes LD enlargement and developmental delay in *C. elegans*. (A) A schematic cartoon for the LD enlargement phenotype under the JUb74 diet. The green dots represent LDs in the intestine. (B) Day-one adults fed with *E. coli* OP50 or the *Microbacterium sp.* JUb74 were supplemented with BODPIY 493/503 to visualize the LDs in live animals. (C) Size distribution of the LDs in wild-type *C. elegans* fed with OP50 or JUb74. Double asterisks indicate *p* < 0.01 in unpaired *t*-test. (D) Two intestinal LD marker strains, LIU1 *ldrIs1[dhs-3p::dhs-3::GFP]* and LIU2 *ldrIs2[mdt-28p::mdt-28::mCherry]*, were fed with JUb74 and imaged at day-one adult stage. The surface of LDs was labelled by fluorescent markers. Scale bar = 20 µm. (E) The percentage of wild-type animals reaching L4 and adult stages when fed with OP50 and JUb74 at 72 and 96 hours after bleaching synchronization. At least fifty animals were used in each experiment with at least two biological replicates. (F) The number of eggs laid by an animal fed with OP50 or JUb74 and the number of eggs that hatched into larvae. At least ten animals were analysed in each condition. Single and double asterisks indicate *p* < 0.05 and *p* < 0.01 in unpaired *t*-tests, respectively. (G) One-cell stage embryos laid by hermaphrodite fed with OP50 or JUb74. Large LD-like structures were observed in the eggs laid by JUb74-fed animals. Scale bar = 20 µm. (H) Species information of *Microbacterium* species that induced large LDs when fed to *C. elegans*. Their 16S rRNA sequences were used to blast the NCBI non-redundant database, and the percentages of identity with known species were shown.

JUb74 was isolated from rotting grapes in Santeuil, France and was previously identified as a strain of the *Microbacterium* genus.^24^ We performed whole genome sequencing of JUb74 and found that it likely belonged to *Microbacterium oleivorans*, which is a Gram-positive and crude-oil degrading *Microbacterium* species.^25^ The sequences of 16S rRNA and 23S rRNA were 99.33% and 98.97% identical to *M. oleivorans*, respectively. JUb74 genome was 2,894,998-bp long and contained 2,776 protein-coding genes (Supplemental Files 1-2).

In addition to promoting the formation of large LDs, the JUb74 diet also caused developmental delay of *C. elegans*. Animals fed with *E. coli* OP50 developed from eggs to adults in three days at 20°C, whereas most animals fed with JUb74 reached adulthood in four days (Figure 1E). We also compared the brood size of the wild-type animals on different diets and found that animals fed with JUb74 laid fewer eggs than the control animals fed with OP50. More importantly, significantly fewer eggs laid by animals fed with JUb74 could hatch, resulting in a much smaller brood size than the animals fed with OP50 (Figure 1F). This small brood size may be caused by the accumulation of large LDs in the embryos (Figure 1G and S1B).

Besides JUb74, three other *Microbacterium* strains in the natural bacteria collection, when fed to *C. elegans*, also induced the formation of enlarged LDs and caused developmental delay and reduced brood size (Figure 1H and Supplemental Files 3-5). We sequenced their 16s rRNA to confirm their genus and identify the most closely related species. Together, the above data indicated that the *Microbacterium* diet altered lipid metabolism, developmental progression, and reproduction in *C. elegans*.

### *Microbacterium* diet is rich in BCFAs and leads to BCFA accumulation in *C. elegans* lipids

*Microbacterium* was known to be rich in BCFAs.^26^ Using gas chromatography-mass spectrometry (GC-MS), we compared the lipid composition of *Microbacterium* JUb74 with *E. coli* OP50 and found drastic differences. 96.5% of the fatty acids in JUb74 were monomethyl BCFAs, of which saturated C15anteiso and C17anteiso were the most abundant lipid species (Figure 2A). In contrast, OP50 mostly contained straight-chain saturated fatty acids (SSFAs; 46%), CFAs (33%), and MUFAs (21%) and had no BCFAs.

**Figure 2.**
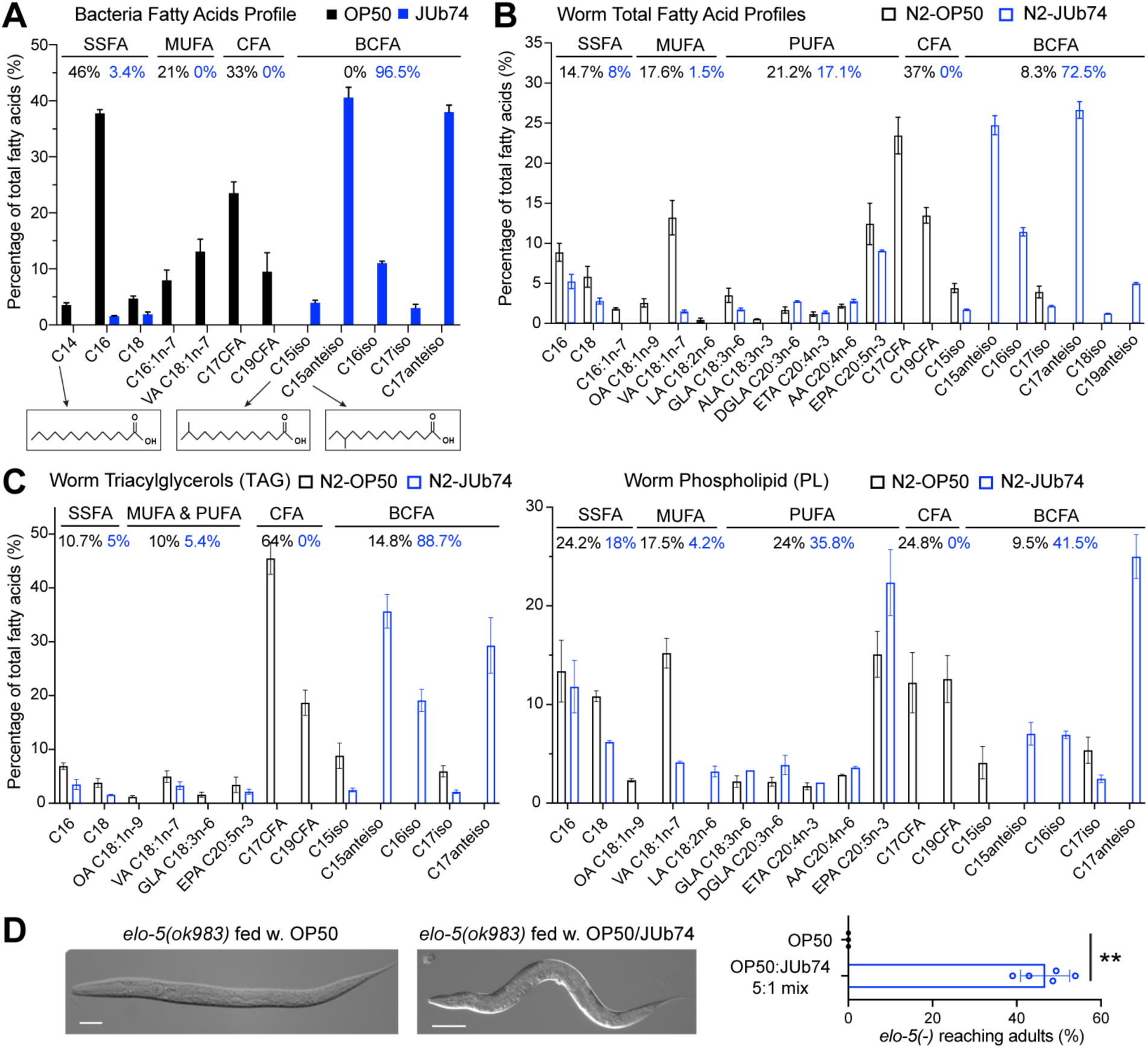
*Microbacterium sp.* JUb74 diet is rich in BCFAs and induces BCFA accumulation in *C. elegans*. (A) GC-MS lipidomic analysis revealed the difference in the total fatty acid profiles of *E. coli* OP50 and *Microbacterium sp.* JUb74. The percentage is defined by the area under the curve for specific fatty acids divided by the area for all detected fatty acids. Combined percentages of SSFAs, MUFAs, CFAs, and BCFAs were shown. Three replicates were performed for each group. The chemical structure of C14, C15iso, and C15anteiso were shown. (B) Total fatty acid profiles of *C. elegans* fed with OP50 and JUb74. (C) Fatty acid profiles of the triacylglycerols (TAG) and phospholipid (PL) fractions of the wild-type *C. elegans* fed with OP50 and JUb74. (D) Images of *elo-5(-)* mutants produced by VC683 *elo-5(ok983) IV/nT1* animals under OP50 diet or a OP50/JUb74 mixed (5:1) diet. Scale bars = 20 µm in the left panel and 100 µm in the right panel. Percentage of *elo-5(-)* mutant animals that reached adulthood under OP50 or OP50/JUb74 mixed diet. Double asterisks indicate *p* < 0.01 in a *t*-test.

We analysed the lipids of animals fed with different bacteria using GC-MS and found that the JUb74 diet significantly altered the lipid composition of *C. elegans*. BCFAs derived from JUb74, including C15anteiso, C16iso, and C17anteiso, were accumulated in *C. elegans* and counted for ∼72% of the total fatty acids (Figure 2B). We also detected C18iso and C19anteiso in animals fed with JUb74, which may be generated by two-carbon elongation from the dietary BCFAs. On the other hand, animals fed with OP50 had mostly SSFAs, CFAs, and MUFAs in their lipids; *de novo* synthesized BCFAs (C15iso and C17iso) counted for only 8.3% of the total lipids.

To understand the BCFA distribution in the triacylglycerol (TAGs) and phospholipids (PL), which make up the LDs and the plasma membrane, respectively, we performed thin-layer chromatography to separate TAG and PL fractions of the total lipids extracted from animals fed with different diets. We found that TAGs in animals fed with JUb74 were made of 88% BCFAs, indicating that the dietary BCFAs were stored in the LDs in the form of TAGs. As a control, TAGs in animals fed with OP50 contained mostly C17 CFA and C19 CFA, and BCFAs made up only 14% of the fatty acids in TAGs (Figure 2C). BCFAs also counted for 41% of the fatty acids in the PL fraction of the lipids in animals fed with JUb74, replacing the abundance of C18:1n-7 *cis*-vaccenic acid and CFAs in animals fed with OP50 (Figure 2C). Our results suggested that the dietary BCFAs can be incorporated into the membrane lipids, although to a lesser extent than storage lipids.

Kniazeva *et al.* previously found that *C. elegans* used the long-chain fatty acid elongases ELO-5 and ELO-6 to *de novo* synthesize C15iso and C17iso, which play essential functions in *C. elegans* development.^27^ As a result, RNAi knockdown of *elo-5* led to arrest at the L1 stage, which could be rescued by dietary supplementation of C17iso. We found that the lethality of the *elo-5* knockout mutation could also be rescued by adding 20% JUb74 to the regular OP50 diet (Figure 2D), suggesting that JUb74-derived BCFAs were biologically functional.

### A high abundance of BCFA in the diet induces enlarged LDs

Next, we ask how JUb74 caused the enlargement of LDs and whether the high abundance (>95% of the total fatty acids) of BCFA in the JUb74 diet was responsible for the enlarged LDs. A *Microbacterium* species (*M. nematophilus*) was found to infect *C. elegans* and colonize the rectal region to cause local swelling.^28,29^ We did not see such phenotype and ruled out that gut colonization was required for the increased LD sizes, since feeding the animals with UV-killed JUb74 still led to the formation of large LDs (Figure 3A). We also found that the innate immunity pathway (e.g., p38 MAPK pathway genes *pmk-1* and *sek-1*)^30^ was not needed for JUb74-induced LD expansion (Figure 3B).

**Figure 3.**
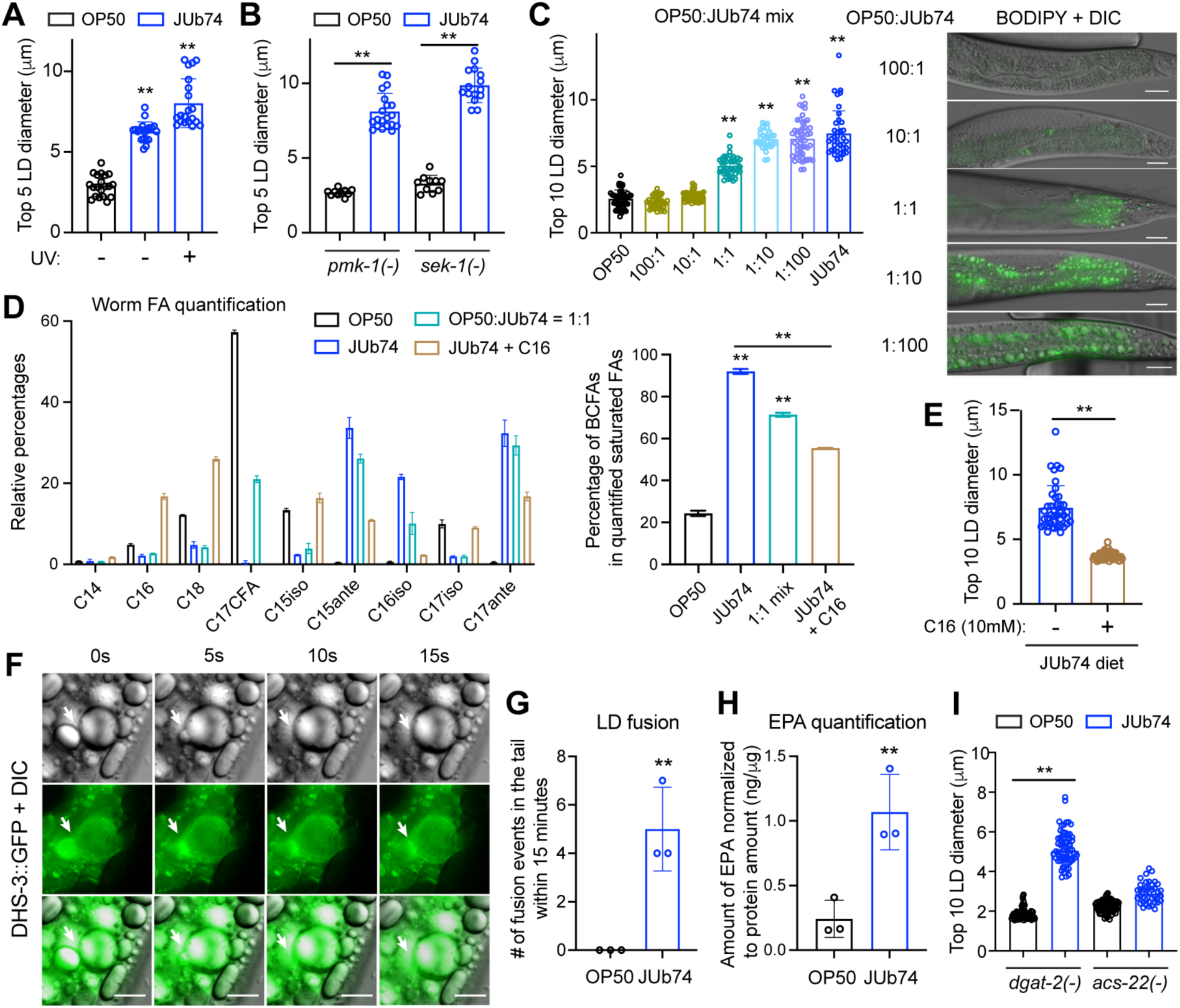
High BCFA abundance in the JUb74 diet induces LD enlargement by promoting LD fusion. (A) The diameter of the five largest LDs (visualized by BODIPY 493/503) in the animals fed with UV-killed OP50 and JUb74. Double asterisks indicate *p* < 0.01 in a Dunnett’s test in comparison with the OP50 diet. (B) The diameters of the five largest LDs in *pmk-1(km25)* and *sek-1(km4)* mutants fed with either OP50 or JUb74. Double asterisks indicate *p* < 0.01 in an unpaired *t*-test. (C) The diameters of the ten largest LDs in animals fed with different ratios of UV-killed OP50 and JUb74. The right panel shows representative images of BODIPY-labelled LDs under different diets. Scale bars = 20 µm. (D) Quantitative GC-MS results for selected SSFAs, CFAs, and BCFAs. The results are normalized as a percentage of the total amount for easy comparison. The bar graph on the right shows the combined percentages of BCFAs among the quantified saturated fatty acids under different diet conditions. Double asterisks indicate *p* < 0.01 in a Tukey’s test in comparison with the OP50 diet or between a specific pair. (E) The diameters of the ten biggest LDs in animals fed with JUb74 grown with or without C16 palmitic acids. (F) Time-lapse images that track the fusion of two LDs in day-one adults of LIU1 *ldrIs1[dhs-3p::dhs-3::GFP]* (GFP labels the LD surface in this strain) fed with JUb74. Arrows point to the interface of two fused LDs. Scale bars = 10 µm. (G) The number of LD fusion events within 15 minutes in the tail region of the animals fed with OP50 or JUb74. (H) GC/MS-quantified EPA amount normalized to total protein concentration in animals fed with OP50 and JUb74. (I) The diameters of the ten biggest LDs in *dgat-2(hj44)* and *acs-22(hj26)* mutants fed with OP50 or JUb74.

We then mixed UV-killed OP50 and JUb74 at different ratios and fed the mixture to *C. elegans* to test how much BCFA was needed in the diet to induce enlarged LDs. We reasoned that if a small amount of certain signaling molecules derived from JUb74 could modulate lipid metabolism in *C. elegans*, we may observe altered LD sizes even when the JUb74 diet was mixed at a low proportion. However, we were only able to observe enlarged LDs at the 1:1 ratio (with BCFAs making up ∼50% of the total fatty acids in the diet) and found that the increase in LD size was dose-dependent on the amount of JUb74 diet or the proportion of BCFA in the diet (Figure 3C and S2A). Only when the OP50-to-JUb74 ratio reached 1:10 and BCFA abundance reached >90%, we were able to observe the same LD size as the animals fed with pure JUb74. These results suggest that the high abundance of dietary BCFA may be responsible for LD enlargement.

To further confirm this idea, we supplemented palmitic acid (C16:0) to the JUb74 culture, which increased the SSFA level to over 60% and reduced the BCFAs to about 35% (Figure S2B), and fed the metabolically modified JUb74 to *C. elegans*. C16 supplementation reduced the LD size to be even smaller than that in the animals fed with the 1:1 mixture of JUb74 and OP50 (Figure 3E), suggesting that the relative abundance of BCFAs controlled the LD size. Using GC-MS, we further quantified selected SSFAs, CFAs, and BCFAs and confirmed the increase of BCFAs in the worm lipid content under the OP50/JUb74 mixed diet compared to the OP50 diet (from ∼20% to ∼70% among the quantified saturated FAs) and the decrease of BCFA percentages (from ∼90% to ∼50%) when palmitic acids were added to the JUb74 diet (Figure 3D). Conversely, we also supplemented either specific BCFAs or a mixture of them to the OP50 culture but were not able to increase the BCFA levels above 30%, likely because OP50 could not take in large amount of BCFAs (Figure S2C). Feeding those BCFA-treated OP50 to wild-type *C. elegans* did not induce large LDs (Figure S2D). Thus, we concluded that, in addition to the dosage effect, a threshold existed for the minimal level of BCFAs that could induce large LDs.

### BCFA-rich lipid droplets are resistant to hydrolysis and prone to fusion

Previous studies of the *C. elegans* mutants that developed enlarged LDs under the regular OP50 diet categorized the mutants into two broad categories, one with defective peroxisomal import or β-oxidation of fatty acid and another with upregulated LD fusion; the former is resistant to hydrolysis, while the latter is not.^19^ We found that the large LDs in animals fed with JUb74 did not shrink or disappear after fasting for 24 or even 72 hours (Figure S3A-B), suggesting that the BCFA-rich LDs were resistant to hydrolysis and could not be easily catabolized. The resistance to hydrolysis may be due to the diet-induced suppression of lipid degradation and/or the physiochemical properties of BCFAs.

Meanwhile, the high frequency of fusion events also led to the growth of large LDs. Using a strain that expressed the LD surface marker DHS-3::GFP, we observed frequent LD fusion in animals fed with JUb74 but not OP50 (Figure 3F-G). In some cases, two medium-sized LDs fused to generate a big one (Figure 3D and Movie S1), while in other cases a big LD absorbed a small one to generate an even bigger LD (Figure S3C and Movie S2). These fusion events appeared to be triggered by the contact between two LDs and occurred most frequently around the vulva and the anus, where the periodic muscle contraction during egg-laying and defecation caused the displacement of LDs and created more chances for them to contact each other.

Three types of LD fusions were previously proposed based on their mechanisms and time scales,^31^ including protein-mediated slow fusion (which takes around 60 minutes), temperature-dependent fusion (which takes 30 seconds), and phospholipid-mediated, proximity-based fast fusion (which occurs within seconds). The LD fusion we observed in *C. elegans* under the BCFA-rich diet occurred in less than 10 seconds, thus fitting the third category. A recent study suggested that PUFAs promote the rapid fusion of LDs in *C. elegans*,^31^ so we hypothesized that increased levels of PUFA may be responsible for the enhanced fusion under the JUb74 diet. Using quantitative GC-MS measurement, we indeed found that animals fed with JUb74 had significantly higher levels of eicosapentaenoic acid (EPA C20:5n-3; the most abundant PUFA in *C. elegans*) than animals fed with OP50 (Figure 3H). Transcriptomic analysis found that animals fed with JUb74 had significantly upregulated expression of the fatty acid desaturase FAT-7 compared to the ones fed with OP50 (see Figure 6 for details), supporting enhanced production of EPA and thus increased LD fusion, which drove the enlargement of LDs.

In addition, we found that the JUb74 diet-induced LD enlargement was dependent on the fatty acyl-CoA synthetase ACS-22/FATP4 but not the diacylglycerol acyltransferase DGAT-2 (Figure 3I). This finding is somewhat surprising since previous studies showed that ACS-22 and DGAT-2 are in the same TAG synthesis complex that promotes LD expansion on the ER-LD interface under the OP50 diet.^14^ We reasoned that DGAT-2 may not mediate the incorporation of branched-chain very-long-chain acyl-CoA into TAGs.

### A gain-of-function mutation in the scavenger receptor *scav-4* enhanced the effects of *Microbacterium* diet on LD enlargement

To understand the genetic pathway that interacts with the *Microbacterium* diet in the regulation of LD size, we performed a forward genetic screen to search for mutants that showed further increased LD size compared to the wild-type animals under the JUb74 diet. In a small-scale screen of a few thousand haploid genomes, we isolated an allele *unk28*, which led to the formation of supersized LDs under the JUb74 diet (Figure 4A). Through chromosomal mapping and whole-genome resequencing, we mapped *unk28* to a missense mutation L462F in *scav-4*, which codes for a CD36 family scavenger receptor.

**Figure 4.**
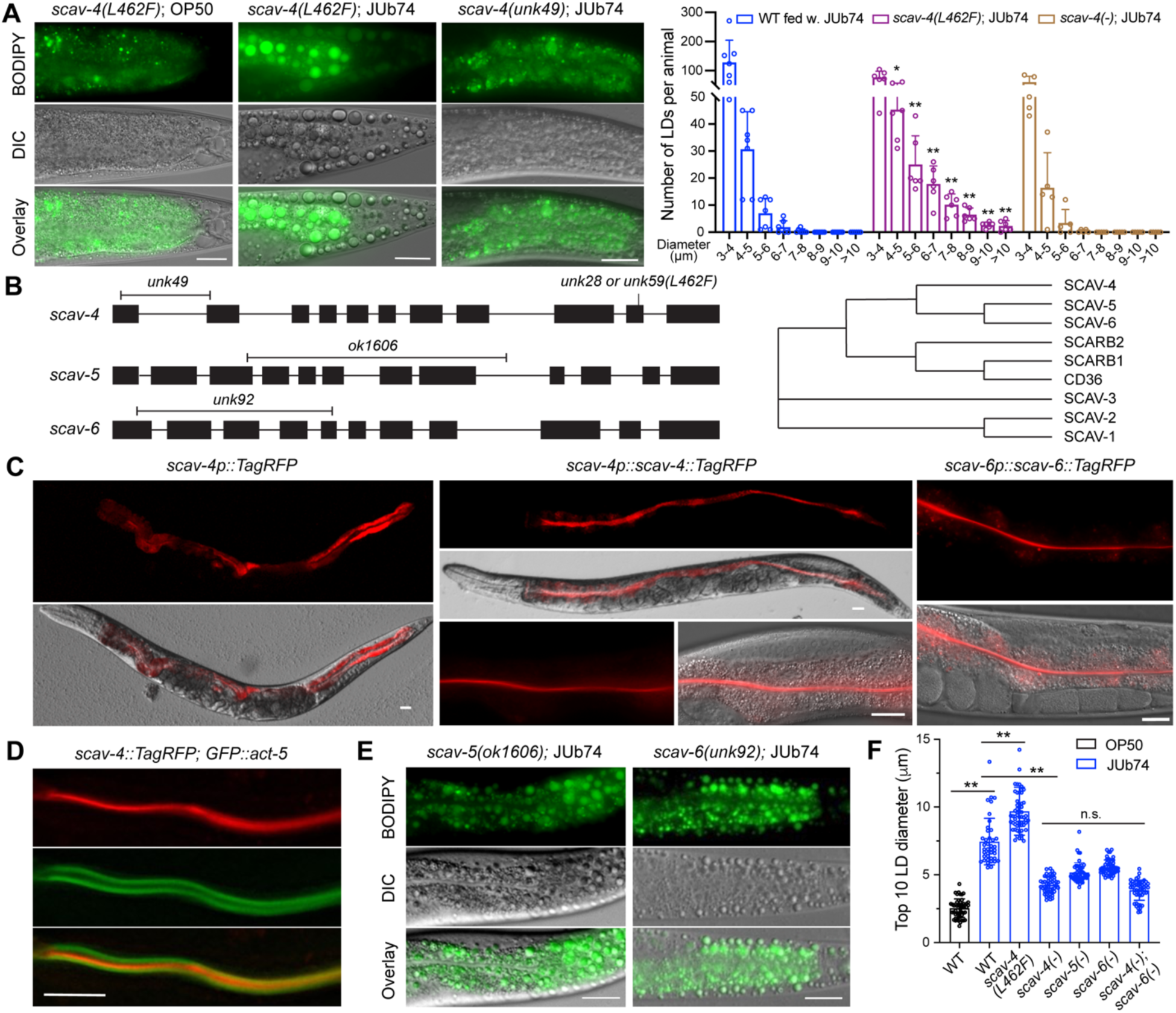
A missense mutation in SCAV-4/CD36 promotes the JUb74-induced LD expansion. (A) BODIPY-stained LDs of *scav-4(unk59; L462F)* and *scav-4* knockout (*unk49*) animals fed with JUb74. Scale bar = 20 µm. LD size distribution of wild-type, *scav-4(unk59)*, and *scav-4(unk49)* animals fed with JUb74. Single and double asterisks indicate *p* < 0.05 and *p* < 0.01, respectively, in a *t*-test comparing the number of LDs in the same size category between the wild type and mutants. (B) Molecular lesions of various *scav-4*, *scav-5*, and *scav-6* alleles and a phylogenetic tree of CD36 scavenger receptor homologs in human (CD36, SCARB1, and SCARB2) and *C. elegans* (SCAV-1∼6). (C) Intestinal expression of the *scav-4p::TagRFP* transcriptional reporter and localization of SCAV-4::TagRFP and SCAV-6::TagRFP translational reporters on the apical membrane of intestinal cells. Scale bar = 20 µm. (D) The close proximity of SCAV-4::TagRFP and GFP::ACT-5 at the apical surface of the intestine. (E) BODIPY-labelled LDs of *scav-5(-)* and *scav-6(-)* knockout mutants fed with JUb74. (F) The diameter of the ten biggest LDs in various strains fed with either OP50 or JUb74. Double asterisks indicate *p* < 0.05 in a post-ANOVA Tukey’s multiple comparison test.

To test whether *scav-4(unk28)* was a loss-of-function allele, we created a *scav-4* knockout allele *unk49* through CRISPR/Cas9-mediated gene editing; *unk49* deleted the first intron and 146 bp in exon 1 and exon 2, causing a frameshift (Figure 4B). Surprisingly, this *scav-4(-)* mutant showed no enlarged LDs when fed with JUb74; instead, they showed even smaller LDs than the wild-type animals under the JUb74 diet (Figure 4A). To confirm *scav-4(L462F)* mutation was indeed the phenotype-causing mutation, we recreated the *scav-4* mutation in the otherwise wild-type background through gene editing, and the resulting *scav-4(unk59)* animals showed the same supersized LDs as the original *scav-4(unk28)* mutants when fed with JUb74. These results suggested that *scav-4(L462F)* is likely a gain-of-function mutation.

JUb74-induced LDs in *scav-4(L462F)* mutants grew to as large as 14 µm in diameter in day-one adults and 31 µm in diameter in day-five adults (Figure S4A and S4B). These supersized LDs were mostly found around the vulva or in the posterior half of the intestine, and their identities as LDs were confirmed by colocalization with intestinal LD markers, DHS-3::GFP (Figure S4C). Importantly, *scav-4(L462F)* mutants did not develop supersized LDs under the OP50 diet, indicating that the effects of this mutation were specific to the BCFA-rich JUb74 diet (Figure 4A and S4D).

### SCAV-4 localizes to the apical membrane of the intestine and may regulate lipid uptake

*C. elegans* genome contains six *scav* genes (*scav-1* to *scav-6*), which are orthologs of the human class B scavenger receptor family proteins, including CD36, SCARB1 (CD36L1 or SR-BI), and SCARB2 (CD36L2 or LIMP-2). These scavenger receptors can bind to lipoproteins and mediate the uptake of lipids and cholesterol into blood cells, liver, adipocytes, and enterocytes.^32,33^ Thus, we hypothesize that SCAV-4 may also function in lipid transport. To analyse the expression pattern of *scav-4* in *C. elegans*, we generated both transcriptional and translational reporters, which were expressed specifically in the intestine (Figure 4C). The SCAV-4::TagRFP fusion proteins were localized to the apical side of the intestine, which is consistent with its role in lipid uptake. In fact, the apical domain marked by GFP::ACT-5 flanked the apical surface labelled by SCAV-4::TagRFP (Figure 4D), further supporting the localization of SCAV-4 on the intestinal membrane facing the lumen. We also noticed that the *scav-4* expression was stronger in the posterior intestine, which was consistent with that enlarged LDs were more concentrated in the posterior half of the intestine under the JUb74 diet.

Since *scav-5* and *scav-6* are paralogs of *scav-4*, we analysed their functions in lipid accumulation using *scav-5(ok1606)* deletion mutants and *scav-6* knockout alleles generated in this study through CRISPR/Cas9-mediated gene editing (Figure 4B). We found that when fed with JUb74, both *scav-5(-)* and *scav-6(-)* mutants had moderately reduced LD sizes, but not to the extent of *scav-4(-)* mutants (Figure 4E). Previous promoter reporter studies showed that *scav-5* and *scav-6* were expressed in the intestine.^34^ We constructed translational reporters for both genes and found weak or no signals for SCAV-5::TagRFP possibly due to low protein levels. The SCAV-6::TagRFP fusion protein was expressed in the intestine and was localized to the apical membrane (Figure 4C). From the fluorescent intensity, the *scav-6* expression appeared to be weaker than the *scav-4* expression. Moreover, *scav-4(-) scav-6(-)* double mutants had the same LD diameter as *scav-4(-)* single mutants (Figure 4F). The above results suggested that SCAV-4 may play a more significant role than the other two paralogs in intestinal lipid uptake.

Supporting this idea, *scav-4(-)* mutants, but not *scav-5(-)* and *scav-6(-)* mutants, showed developmental delay and smaller brood size and had thinner bodies compared to the wild-type animals even under the OP50 diet (Figure 5C and 5D) likely due to defects in lipid absorption. In fact, GC-MS analysis showed that *scav-4(-)* mutants had reduced percentages of CFA (obtained from the bacterial diet) and increased percentages of PUFA (*de novo* synthesized) compared to the wild-type animals (Figure S5A), supporting that SCAV-4 functions as a fatty acid importer.

**Figure 5.**
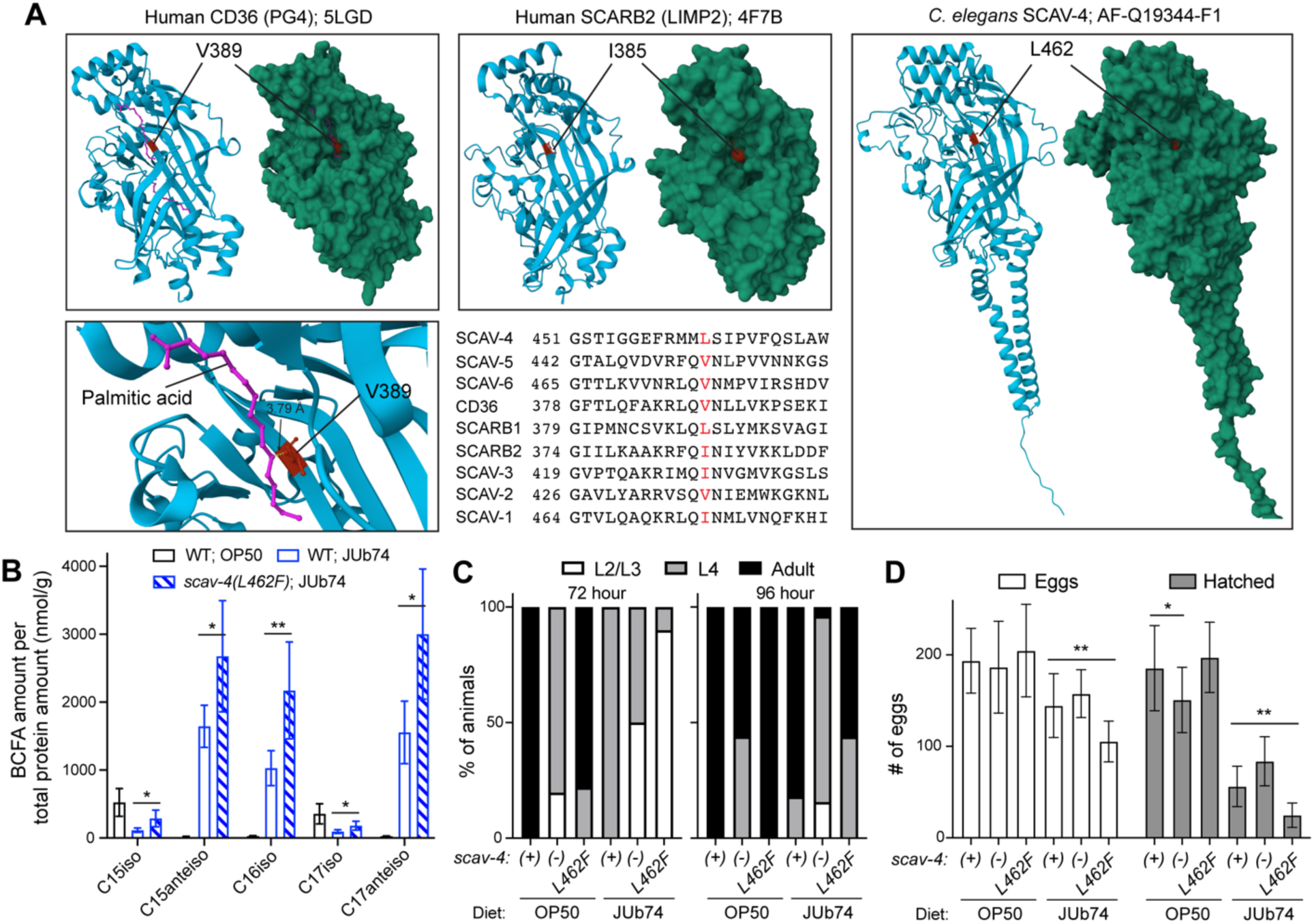
SCAV-4(L462F) mutation affects BCFA absorption, development, and reproduction. (A) Location of the residues homologous to SCAV-4 L462 in the lipid-conducting tunnel of CD36 and SCRAB2 (LIMP2) based on the reported protein structures in PDB (5LGD and 4F7B, respectively) and the location of the L462 in an Alphafold-predicted SCAV-4 structure (AF-Q19344-F1). The CD36 structure is bound to palmitic acid (pink) and the Valine 389 residue is in contact distance (3.79Å) from the palmitic acid molecule (lower left panel). (B) Mass spectrometry-based quantification of BCFAs in wild-type and *scav-4(L462F)* animals fed with JUb74. Single and double asterisks indicate *p* < 0.05 and *p* < 0.01 in a *t*-test, respectively. (C) The percentage of animals reaching L2/L3, L4, and adult stages in wild-type [*scav-4(+)*], *scav-4(unk49)* [*(-)*], and *scav-4(unk59; L462F)* animals fed with OP50 or JUb74 at 72 hours and 96 hours after bleaching. (D) The number of eggs laid and hatched per animal that was fed with OP50 or JUb74. Double asterisks indicate *p* < 0.01 in an unpaired *t*-test.

### L462F mutations in SCAV-4 may enhance BCFA uptake

Since the L462F mutation in *scav-4* led to supersized LDs under the BCFA-rich JUb74 diet, we next mapped the mutation onto the sequence and structure of CD36 scavenger receptors. The L462 residue in SCAV-4 appeared to be conserved among the *C. elegans* SCAV proteins and human CD36 scavenger receptors and is homologous to V389 in human CD36, L390 in SCARB1, and I385 in SCARB2 (Figure 5A). Based on the solved structure of palmitic acid-bound CD36,^35^ we found that the homologous V389 residue is located in the lipid conducting tunnel close to the entrance and is within contact distance (3.79 Å) with the lipid molecule for hydrophobic interactions (Figure 5A). Similarly, the I385 residue in a crystal structure of human LIMP2/SCARB2 and the L462 residue in an AlphaFold-predicted structure of *C. elegans* SCAV-4 also appeared to be located in the lumen of the cavities that presumably form the lipid tunnel (Figure 5A). These results suggest that the L462F mutation may affect the transport of BCFAs under the JUb74 diet, which could possibly lead to supersized LDs.

To measure BCFA absorption, we quantified the amount of various BCFA species in *scav-4* mutants fed with the JUb74 diet using GC-MS. We found that *scav-4(L462F)* mutants indeed accumulated more C15iso, C15anteiso, C16iso, C17iso, and C17anteiso than the wild-type animals (Figure 5B), confirming that the SCAV-4 L462F mutation enhanced BCFA absorption in the intestine. The overall lipid profiles of *scav-4(L462F)* mutants also showed moderately higher percentages of BCFAs than the wild-type animals when fed with JUb74, while *scav-4(-)* mutants had significantly lower BCFA percentages (Figure S5B), supporting the increased and decreased BCFA uptake in the two mutants, respectively.

Given that the BCFA-rich diet induced developmental delay and reduction in brood size, we compared the response of wild-type animals and *scav-4* mutants to JUb74 and found that *scav-4(L462F)* mutants had a more severe phenotype than the wild-type animals under JUb74 diet, although they did not show much difference under the OP50 diet (Figure 5C-D). We reasoned that the enhanced BCFA absorption in *scav-4(L462F)* mutants may exacerbate the detrimental effects of BCFA accumulation on the development and reproduction of the animals. Interestingly, although *scav-4(-)* animals showed slow development in general due to lipid uptake defects, their response to JUb74 diet was weaker than the *scav-4(L462F)* mutants, further supporting that the missense mutation in *scav-4* has gain-of-function effects on the animals’ response to BCFA-rich diet.

### Mutations in *scav-4* altered the transcriptomic response to the JUb74 diet

To understand how mutations in *scav-4* modulate the interaction between the JUb74 diet and the host, we conducted transcriptomic analyses on wild-type, *scav-4(L462F)*, and *scav-4(-)* animals fed with OP50 and JUb74 through RNA sequencing. Using log2(fold change) above 2 or below -2 as the threshold (adjusted *p* value < 0.05), we found that only 4 genes showed differential expression between the wild-type and *scav-4(L462F)* under the OP50 diet, whereas 3,191 genes were upregulated and 1,053 genes were downregulated in *scav-4(L462F)* mutants compared to the wild type under the JUb74 diet (Figure 6A; Table S1-S2). This finding supported that *scav-4(L462F)* did not alter lipid metabolism under the regular OP50 diet, and the L462F mutation in SCAV-4 specifically affected how the animal responded to the BCFA-rich JUb74 diet. In contrast, we found 1,942 upregulated and 310 downregulated genes in *scav-4(-)* mutants compared to the wild type when both were fed with OP50, suggesting that SCAV-4 plays a physiological role under the regular OP50 diet (Table S3-S4). Nevertheless, under the JUb74 diet, more extensive changes in the transcriptomic profiles were observed between the wild-type and *scav-4(-)* animals (Figure 6A).

**Figure 6.**
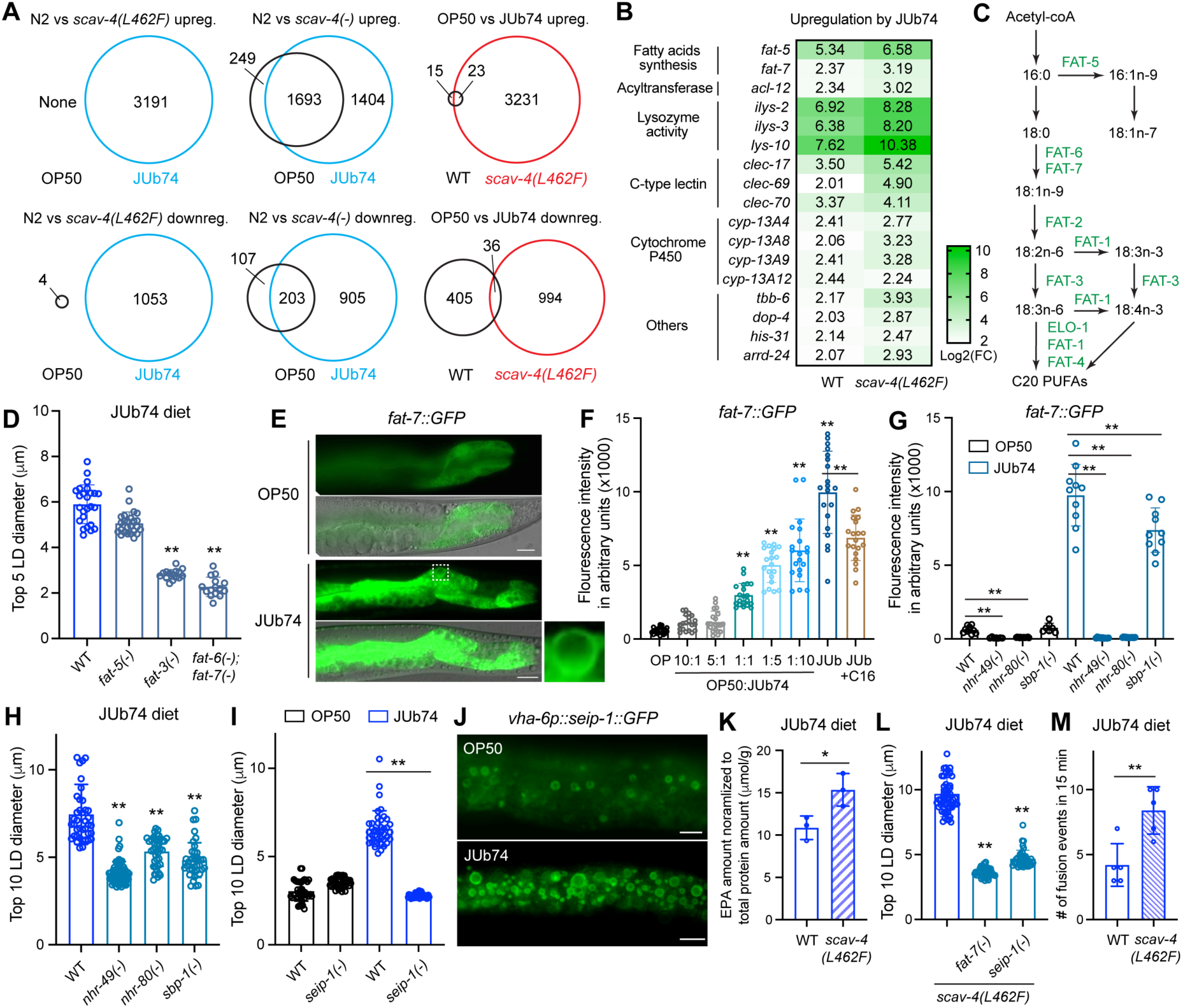
JUb74 diet induces the upregulation of the desaturase *fat-7* to promote LD expansion in a Seipin-dependent manner. (A) The left four Venn diagrams show the overlap of significantly upregulated or downregulated genes in *scav-4(unk59; L462F)* and *scav-4(unk49)* mutants compared to the wild-type animals in OP50 and JUb74 diet. The right two Venn diagrams show the overlap of upregulated or downregulated genes under the JUb74 diet compared to the OP50 diet in the wild-type and *scav-4(L462F)* animals. (B) Selected genes among the 23 genes commonly upregulated by the JUb74 diet in both wild-type and *scav-4(L462F)* animals. The log2(fold change) of the upregulation is shown in a heatmap. (C) The genetic pathway that controls the synthesis of MUFAs and PUFAs in *C. elegans*. The desaturases that control each step are shown. (D) The diameters of the five biggest LDs in *fat-3(ok1126)*, *fat-5(tm420)*, and *fat-6(tm331); fat-7(wa36)* mutants fed with JUb74. Double asterisks indicate *p* < 0.01 in a Dunnett’s test in comparison with the wild type. (E) The upregulation of *waEx15[fat-7::GFP]* under JUb74 diet compared to the OP50 diet. The dashed box is enlarged to show the enriched FAT-7::GFP signal surrounding the enlarged LD. Scale bar = 20 µm. (F) Fluorescent intensity of *fat-7::GFP* expressed from the *nIs590[fat-7p::fat-7::GFP]* transgene was quantified under various diets; representative images are in Figure S7A-B. Double asterisks indicate *p* < 0.01 in a Tukey’s test in comparison with the OP50 diet or between specific pairs. (G) Fluorescent intensity of *fat-7::GFP* expressed from *nIs590* in *nhr-49(nr2041)*, *nhr-80(tm1011)*, and *sbp-1(ep79)* mutants under OP50 or JUb74 diet. (H-I) Diameter of the ten biggest LDs in various mutants fed with OP50 or JUb74. (J) Enrichment of *seip-1::GFP* expressed from *hjSi3[vha-6p::seip-1::GFP]* under OP50 and JUb74 diet. (K) The amount of EPA in wild-type animals and *scav-4(L462F)* mutants fed with JUb74 normalized to the total protein content of the animal. (L) Diameters of the ten biggest LDs in *scav-4(unk59)* double mutants with *fat-7(-)* or *seip-1(-)* mutants fed with JUb74. (M) The number of fusion events observed in the tail region within 15 minutes in wild-type and *scav-4(unk59)* adults fed with JUb74.

Unexpectedly, the same set of genes was altered in both *scav-4(L462F)* and *scav-4(-)* mutants compared with the wild type when fed with JUb74. The upregulated genes were enriched in phosphorylation and signal transduction, while the downregulated genes were enriched in organismal development, morphogenesis, and transcription (Figure S6). We reasoned that these transcriptomic changes were related to the slower development observed in both gain-of-function and loss-of-function *scav-4* mutants compared to the wild-type animals under the JUb74 diet (Figure 5C-D). An optimal level of SCAV-4 activity helps the animal adapt to the BCFA-rich diet.

We next analysed the gene expression changes induced by the JUb74 diet. In the wild-type animals, the JUb74 diet upregulated 38 genes and downregulated 441 genes compared to the OP50 diet (Table S5); the effects of the JUb74 diet were much stronger in *scav-4(L462F)* mutants, causing 3,254 and 1,030 genes to be upregulated and downregulated, respectively (Figure 6A; Table S6). This exaggerated transcriptomic response to the JUb74 diet was consistent with stronger phenotypes in LD enlargement, developmental delay, and brood size reduction in *scav-4(L462F)* mutants (Figure 5C-D). Among the 23 genes commonly upregulated in both wild-type and *scav-4(L462F)* animals by JUb74 diet, we found three lysozymes (*lys-10*, *ilys-2*, and *ilys-3*), five innate immune genes (*clec-17*, *clec-69*, *clec-70*, *Y38H6C.23*, and *F01D5.3*), four cytochrome P450 genes (*cyp-13A4, cyp-13A8, cyp-13A9,* and *cyp-13A12*), two fatty acid desaturases (*fat-5* and *fat-7*), one acyltransferase (*acl-12*), and one carnitine O-palmitoyltransferase (*W03F9.4*) (Figure 6B). The first two groups of genes may be involved in bacterial recognition, and the other genes are involved in lipid metabolism. Most of these JUb74-induced genes had bigger expression changes in *scav-4(L462F)* mutants than the wild-type animals (Figure 6B), which is consistent with a stronger metabolic response in the mutants.

### JUb74-induced LD enlargement depends on fatty acid desaturases

The finding of JUb74-induced upregulation of *fat-5* and *fat-7* was particularly interesting, because FAT-5 and FAT-7 catalyse Δ9-desaturation for C16 and C18 fatty acids, respectively (Figure 6C). Their upregulation could lead to elevated amounts of *de novo* synthesized MUFA and PUFAs. To test whether the upregulation of *fat-5* and *fat-7* under JUb74 diet contributed to the enlargement of LDs, we examined the mutants of the desaturases and found that *fat-6(-) fat-7(-)* double mutants, in which the production of C18:1n-9 and the subsequent C18 and C20 PUFAs were blocked, failed to develop enlarged LDs under JUb74 diet (Figure 6D). Furthermore, mutations in *fat-3*, which codes for a Δ6-desaturase essential for the generation of C20 PUFAs, also led to the failure of developing enlarged LDs. In contrast, the *fat-5(-)* mutants, in which only the synthesis of C16:1n-9 and C18:1n-7 MUFAs were blocked, were able to generate large LDs under the JUb74 diet (Figure 6D). The above results suggested that PUFAs and not MUFAs were responsible for the LD expansion. The most abundant PUFA in *C. elegans* is EPA, and we observed an increased amount of EPA in animals fed with JUb74 (Figure 3H), suggesting that the upregulation of *fat-7* may lead to the increased production of EPA, which then causes LD enlargement by promoting LD budding and growth from the endoplasmic reticulum^15^ and LD fusion.^31^

Using two independent *fat-7* translational reporters, we confirmed the upregulation of *fat-7* by JUb74 (Figure 6E and S7A). Strikingly, FAT-7::GFP signal appeared to be enriched in a ring-like shape surrounding the enlarged LDs under the JUb74 diet, while it was mostly diffusive in the intestine under the OP50 diet (Figure 6E and S7A). The significance of this localization pattern is unclear, but we suspect that it may facilitate the accumulation of PUFA on the LD surface. Furthermore, we found that the level of *fat-7* was regulated by the abundance of BCFA in the diet. When we fed the reporter strain with a mixture of OP50 and JUb74, the *fat-7::GFP* only started to show upregulation when the JUb74 ratio was above 50% (Figure 6F), similar to the threshold for the large LD phenotype (Figure 3C). The level of *fat-7* appeared to be dose-dependent on the amount of JUb74. Moreover, when we supplemented C16 to reduce the amount of BCFAs in the JUb74 diet, we observed significantly reduced *fat-7::GFP* levels, further supporting that the BCFA abundance regulates *fat-7* expression (Figure 6F).

Since *fat-7* expression is known to be regulated by the nuclear hormone receptors NHR-49^36^ and NHR-80,^37^ which form a heterodimer,^38^ and the SREBP homolog *sbp-1*,^39^ we tested whether they were involved in BCFA-induced upregulation of *fat-7*. Mutations in either *nhr-49* and *nhr-80* abolished the upregulation of *fat-7::GFP* by the JUb74 diet and prevented the LD enlargement (Figure 6G, 6H, and S7B), suggesting that the metabolic regulator NHR-49/NHR-80 responded to the JUb74 diet and mediated the activation of *fat-7* expression. On the other hand, the loss of *sbp-1* only slightly reduced *fat-7* expression but still reduced LD size, suggesting that other SBP-1 downstream genes may be required for JUb74-induced LD enlargement.

### PUFAs promote LD enlargement by enhancing SEIP-1 recruitment and LD fusion

PUFAs promote LD budding and growth by recruiting SEIP-1/seipin into an ER subdomain that is associated with nascent LDs.^15^ We found that the *seip-1(-)* mutants did not form enlarged LDs when fed with JUb74, suggesting that the recruitment of SEIP-1 by PUFA was required for LD expansion under the BCFA-rich diet (Figure 6I). In fact, we observed the enrichment of SEIP-1::GFP on the periphery of the enlarged LDs in animals fed with JUb74 (Figure 6J), supporting that JUb74 diet may indeed enhance LD budding from the ER in a SEIP-1-dependent manner.

We next tested whether the formation of supersized LDs in the *scav-4(L462F)* mutants was caused by not only enhanced absorption of BCFAs but also increased production of PUFAs. Using quantitative GC-MS, we first found that *scav-4(L462F)* mutants had stronger upregulation of *fat-7* (Figure 6B) and produced higher amount of EPA than the wild-type animals when both were fed with JUb74 (Figure 6K). This increased amount of PUFAs was responsible for the larger LD sizes in *scav-4(L462F)* mutants compared to the wild type, because mutations in *fat-7* blocked the production of PUFAs and prevented the generation of supersized LDs in *scav-4(L462F)* mutants (Figure 6L). The supersized LDs also depended on *seip-1* in *scav-4(L462F)* mutants (Figure 6L), and the mutants had a higher LD fusion rate than the wild-type animals (Figure 6M). In one case, we observed five LDs fuse into a big one within 2 minutes, highlighting the high frequency of LD fusion in the mutants (Figure S7C and Movie S3). Thus, the higher level of PUFAs in *scav-4(L462F)* mutants may aggravate LD enlargement by further stimulating the budding and fusion of LDs.

### BCFAs promote lipogenesis by repressing AMPK and activating mitoUPR

We next focused on the genes downregulated by the JUb74 diet in both wild-type and *scav-4(L462F)* mutants. The top downregulated gene *argk-1* codes for a creatine kinase that is repressed by the S6 kinase (S6K) RSKS-1 and acts upstream of the energy sensor AAK-2/AMP-activated protein kinase (AMPK),^40^ which is known to repress fatty acid synthesis and activate β-oxidation in response to low energy levels^41^ (Figure 7A-B). Interestingly, recent studies found that monomethyl BCFAs trigger the mTORC1 pathway,^42^ which can activate RSKS-1/S6K,^43^ which in turn inhibits ARGK-1 and AMPK.^40^ Thus, the BCFA-rich JUb74 diet may induce large LDs by inhibiting the AMPK pathway.

**Figure 7.**
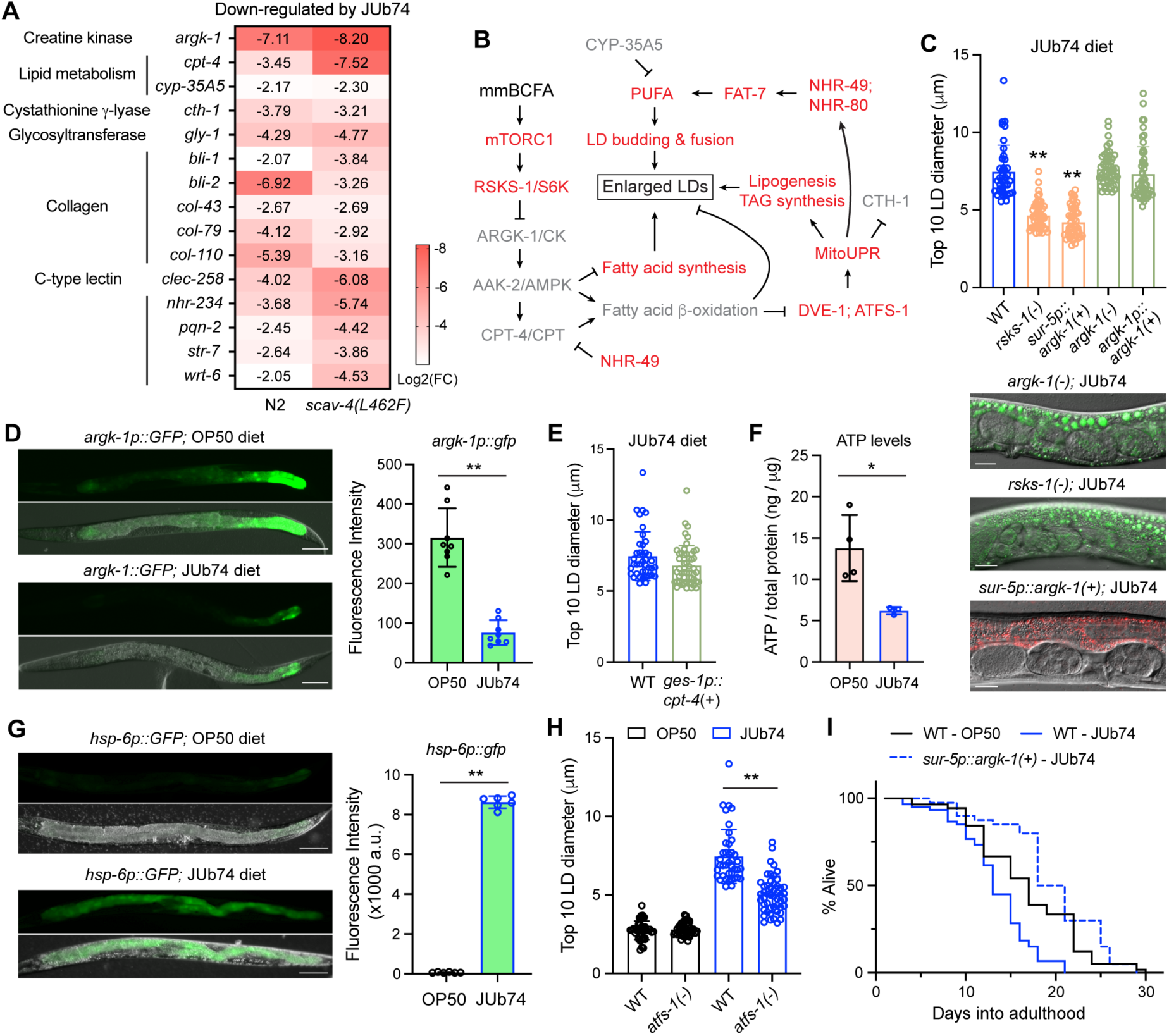
BCFAs inhibit *argk-1* expression and the AMPK pathway and activate mitoUPR to promote lipogenesis. (A) Selected genes among the 36 genes commonly downregulated by the JUb74 diet in both wild-type and *scav-4(L462F)* animals. The log2(fold change) of the downregulation are shown in a heatmap. (B) The signaling network activated by monomethyl BCFAs. Genes and pathways in red are activated by BCFAs, while items in grey are inhibited by high levels of BCFAs. (C) Diameters of the ten biggest LDs in wild-type, *rsks-1(ok1255)*, *sqIs32[sur-5p::argk-1::GFP]*, *argk-1(ok2993)*, and *sqEx9[argk-1p::argk-1::mCherry]* animals. Representative images were shown. Scale bar = 20 µm. (D) Expression of *sqEx82[argk-1p::GFP]* under OP50 and JUb74 diet. Scale bar = 100 µm. Quantification of GFP intensity is shown on the right. Double asterisks indicate *p* < 0.01 in a *t*-test. (E) Diameters of the ten biggest LDs in animals carrying the *unkEx732[ges-1p::cpt-4]* transgene. No significant differences were found in comparison with the wild type. (F) ATP levels of wild-type animals fed with OP50 or JUb74, normalized to total protein amount. (G) The expression of mitoUPR marker *hsp-6p::GFP* under OP50 or JUb74 diet. (H) LD size in wild-type and *atfs-1(gk3094)* animals fed with OP50 and JUb74 diet. (I) Survival curve of wild-type and *sqIs32[sur-5p::argk-1::GFP]* animals fed with OP50 or JUb74. Statistical significance (*p* < 0.01) was found between WT-OP50 and WT-JUb74 groups and between WT-JUb74 and *sur-5p::argk-1(+)*-JUb74 groups in a log-rank (Mantel-Cox) test.

To confirm this idea, we found that *rsks-1(-)* mutants had significantly reduced LD sizes than wild-type animals under the JUb74 diet (Figure 7C) likely due to the upregulation of ARGK-1 levels in *rsks-1(-)* mutants.^40^ Overexpression of *argk-1* from a heterologous promoter also prevented the formation of enlarged LDs (Figure 7C), supporting that ARGK-1 is a key regulator whose repression led to lipid accumulation and LD growth under the JUb74 diet. Interestingly, transgenic *argk-1* overexpression from its own promoter did not restore the normal LD size (Figure 7C), likely because transcription from the *argk-1* promoter was repressed by the BCFA-rich diet. In fact, the expression of *argk-1p::GFP* reporter was strongly inhibited in the JUb74 diet compared to that under the OP50 diet (Figure 7D).

AMPK promotes lipid catabolism by inhibiting acetyl-CoA carboxylase 2 (ACC2), which generates malonyl-CoA that inhibits carnitine palmitoyltransferase 1 (CPT1), an essential enzyme in β-oxidation.^41^ Among the JUb74-downregulated genes, *cpt-4* codes for a *C. elegans* homolog of CPT; its repression at the transcriptional level (possibly by *nhr-49*^44^) may further inhibit fatty acid oxidation in addition to the functional inhibition by low AMPK activity. However, overexpression of *cpt-4* was not sufficient to reduce LD size under the JUb74 diet (Figure 7E), suggesting that the key regulatory point of the signaling network was likely upstream of CPT-4 and at the control of AAK-2 activity by ARGK-1. The inhibition of AMPK pathway generally leads to reduced energy production; indeed, we found that the ATP levels in animals fed with JUb74 were significantly lower than the animals fed with OP50 (Figure 7F). AMPK inhibition rendered the animals insensitive to low ATP levels, and this insensitivity prevented fatty acid oxidation and exacerbated energy deprivation.

Previous studies found that pharmacological inhibition of CPT1 triggers mitochondrial unfolded protein response (mitoUPR), which then activates lipogenesis and TAG synthesis genes to cause lipid accumulation.^45,46^ We tested whether mitoUPR contributed to the enhanced lipid biosynthesis under JUb74 diet. First, we found that the mitoUPR indicator *hsp-6p::GFP* was strongly activated when the animals were fed with JUb74 (Figure 7G); second, the cystathionine γ-lyase *cth-1* was known to be repressed by mitoUPR^47^ and was found among the JUb74-downregulated genes; third, mitoUPR activates the expression of acyltransferase *acl-12*,^45^ which was upregulated by the JUb74 diet (Figure 6B); fourth, mitoUPR is mediated by the transcription factor DVE-1 and ATFS-1; deleting *atfs-1* reduced the LD sizes under JUb74 diet (Figure7H), confirming that the activation of mitoUPR coordinates the inhibition of fatty acid degradation and the promotion of lipid synthesis for LD enlargement. Finally, mitoUPR is known to activate *nhr-80*,^46^ which is required for the JUb74-induced upregulation of *fat-7*, thus connecting mitoUPR with the enhanced production of PUFA (Figure 7B). Supporting this signaling regulation, we observed increased NHR-80 nuclear accumulation under the JUb74 diet (Figure S7D).

### Overexpression of *argk-1* rescues the shortened lifespan under a BCFA-rich diet

The *rsks-1/argk-1/aak-2* pathway is known to regulate lifespan, since *rsks-1(-)* mutants showed extended lifespan due to the accumulation of ARGK-1^40^ and the loss of AAK-2/AMPK shortened lifespan.^48^ In addition, knocking down *cth-1* caused a mild decrease in lifespan,^49^ and the nuclear receptor *nhr-234* and the cytochrome P450 *cyp-35A5* were required for remofuscin-induced longevity;^50^ all three genes are downregulated by the JUb74 diet (Figure 7A). Thus, we predicted that the JUb74 diet may reduce the lifespan of *C. elegans*. Indeed, we found that feeding with JUb74 significantly shortened the lifespan of the wild-type animals, while overexpression of *argk-1* was able to reverse the JUb74 diet-induced shortening of lifespan (Figure 7I). Interestingly, overexpression of *argk-1* was able to elongate lifespan in both OP50 diet^40^ and JUb74 diet (this study), suggesting a general role of ARGK-1 in promoting longevity regardless of the lipid composition in the diet. The lifespan of *scav-4(L462F)* mutants were similarly affected by the JUb74 diet and did not show a significant difference from the wild-type animals under both diets, suggesting that the formation of supersized LD size did not further shorten lifespan (Figure S7E).

## DISCUSSION

### BCFA abundance in the diet promotes lipid storage and inhibits energy consumption

Monomethyl BCFAs were recently found to mediate amino acid sensing by activating mTORC1 in *C. elegans* and mammalian adipocytes.^42^ Specifically, leucine-derived or dietary C17iso was used to synthesize glucosylceramides, which is critical for the activation of the mTORC1 pathway.^42^ In a mouse model, dietary proteins from a mixed source exacerbated high-fat diet-induced obesity by promoting the mTORC1/S6K signaling and by elevating microbial production of BCFAs in the gut,^51^ suggesting a second mechanism by which BCFAs can act as a signal downstream of amino acid sensing. In this study, we found that when the abundance of dietary BCFAs reaches a certain threshold, they can affect the homeostasis of lipid metabolism through the mTORC1/S6K/AMPK pathway. The overactivation of mTORC1 by high levels of BCFAs led to the downregulation of ARGK-1/CK and strong inhibition of AAK-2/AMPK pathway, which resulted in reduced fatty acid β-oxidation and increased lipogenesis and TAG synthesis. The consequence is that animals make large LDs to store lipids instead of degrading them while under energy stress. Thus, abundant BCFAs in the diet appear to trick the animals into sensing that their energy level is high and therefore inhibit the mobilization of fatty acids for energy production. Similarly, dietary BCFAs can help bypass the amino acid-sensing checkpoint and force the continuation of development under amino acid deficiency.^42^ In this sense, dietary BCFAs can disable the host mechanisms of nutrient sensing.

Another interesting finding in this study is the connection between dietary BCFAs and increased production of PUFA through mitoUPR. Previous studies showed that reduced fatty acid oxidation and increased lipid accumulation promote nuclear translocation of DVE-1 and ATFS-1,^45^ which not only activate genes related to mitoUPR but also upregulate the metabolic regulator *nhr-80*,^46^ which in turn activates the diacylglycerol acyltransferase *dgat-2* to promote TAG synthesis^46^ and upregulates fatty acid desaturases *fat-5* and *fat-7* to promote PUFA production.^37^ Elevated PUFA levels facilitate LD budding, expansion, and fusion, so that the enlarged LDs can accommodate the excessive TAGs generated through enhanced lipogenesis. We found that dietary BCFAs turns on this signaling network to upregulate *fat-7* expression in a dose-dependent manner that mirrors their effects on LD expansion, suggesting that PUFA is essential in BCFA-induced lipid accumulation.

Besides the signaling role, whether BCFAs have other properties that are not shared by SSFAs is unclear. The enzymes involved in fatty acid synthesis and degradation may have different specificities or efficiencies towards BCFAs and SSFAs. For example, the 3-ketoacyl-acyl carrier protein (ACP) synthase III (FabH) in *E. coli* has a strong preference for acetyl-CoA as a primer, so *E. coli* only make SSFAs;^52^ while FabH proteins in many Gram-positive bacteria are highly selective in using isovaleryl-CoA, isobutyryl-CoA, and 2-methylbutyryl-CoA as substrate, so they mostly produce BCFAs.^53^ By the same token, the enzymes involved in β-oxidation (e.g., acyl-CoA dehydrogenase, enoyl-CoA hydratase, hydroxy acyl-CoA dehydrogenase, and ketoacyl-CoA thiolase) might also have different preferences towards SSFAs and BCFAs, making them inequivalent in energy production. Further studies are needed to test whether BCFA-rich TAGs or SSFA-rich TAGs are easier to catabolize by the same enzymes.

### Detrimental effects of high-BCFA diet on development, reproduction, and lifespan

Despite comprehensive studies of lipid metabolism in the past, the health effects of dietary BCFAs, which count for 2% of dairy fat, are understudied. Nutritional studies found that BCFA levels were inversely correlated with body mass index,^54^ and that BCFA levels were higher in the adipose tissue of lean people compared to obese individuals and were associated with insulin sensitivity,^4^ suggesting potentially beneficial effects of BCFAs in weight control. BCFAs have also shown anti-inflammatory^9,55^ and anticarcinogenic effects^10^ in cell culture studies. However, studies directly assessing the metabolic effects of dietary BCFAs in humans or rodents are still lacking.

In this study, we found that BCFA levels in the dietary fat above a certain threshold led to excessive lipid accumulation, developmental delay, impaired reproduction, and shortened lifespan, suggesting that a high-BCFA diet may also have detrimental effects likely due to its repression of ARGK-1/CK and inhibition of the AMPK pathway. Although the BCFA content in the human diet may never reach the total lipid percentages used in our experiments, BCFAs can also be synthesized from branched chain amino acids (BCAAs) by human cells and be produced by the gut microbiota. Thus, it is possible that the amount of BCFAs in the human body might exceed a certain threshold (e.g., under some pathological conditions) to cause detrimental effects. Understanding the mechanisms underlying the health impacts of high BCFA levels can help prevent such potential harms.

### Genetic variation in SCAV-4/CD36 influences host susceptibility to high-BCFA diet-induced metabolic syndrome

Through genetic studies, we isolated a *scav-4* missense mutation (which alters an evolutionary conserved amino acid residue) that increased BCFA absorption and enhanced the metabolic phenotypes induced by a high-BCFA diet. Since *scav-4* is a homolog of human CD36, our findings suggest that genetic variations in this fatty acid transporter can predispose certain individuals to the health impact of dietary BCFA. Interestingly, the *scav-4(L462F)* mutants grew normally as the wild-type animals under the regular OP50 diet that did not contain BCFA, and there was little transcriptomic difference between the mutants and the wild type under the OP50 diet. Thus, the L462F variation has a diet-specific phenotype, and its deleterious effect can be prevented if the high-BCFA diet is avoided.

In humans, many variants in CD36 are associated with dietary fat intake and high serum TAG levels,^56^ as well as increased risks of obesity, cardiovascular disease, and type II diabetes.^57,58^ However, no CD36 variants were reported to respond to specific types of dietary lipids yet, likely because such variants would only manifest their effects under specific diets and are entirely neutral under other diet conditions. L462 in SCAV-4 is homologous to V389 in CD36. Although variants affecting V389 were not reported before, an R386W variant in the NCBI ClinVar database affected a residue on the same β-strand as V389. This variant has a 0.29% frequency in an East Asian population and is classified as “likely pathogenic” for platelet glycoprotein IV (CD36) deficiency, suggesting that it may be a loss-of-function allele.^59^ Whether any CD36 polymorphism in humans specifically reduces or enhances BCFA uptake and thus changes the host susceptibility to BCFA-rich diet-induced metabolic syndromes awaits further studies.

## MATERIALS AND METHODS

### Strains, DNA constructs, and transgenes

*C. elegans* wild-type (N2) and mutant or transgenic strains were maintained at 20°C using standard culturing protocol^60^. A previously published collection of bacteria found in the natural habitat of *C. elegans* was kindly provided by Dr. Gary Ruvkun^24^. Some strains were provided by the *Caenorhabditis* Genetics Centre (CGC), which is funded by the National Institutes of Health Office of Research Infrastructure Programs (P40 OD010440), or by the National BioResource Project supported by the Japanese government. The following alleles and transgenes were generous gifts from Dr. Ho Yi Mak, Hong Kong University of Science and Technology: *dgat-2(hj44) V*, *seip-1(tm4221) V, acs-22(hj26) X*, *hjSi62[mRuby::dgat-2]*, *hjSi3[vha-6p::seip-1 cDNA::GFP_TEV_3xFLAG::let-858 3′UTR] II.* The *scav-4(unk28*; L462F) was isolated from a forward genetic screen searching for mutants that affect LD growth under the JUb74 diet and were mapped through outcross and whole-genome resequencing. All strains used in this study are listed in Table S7.

To create transcriptional and translational reporters for *scav-4*, we cloned a 1.9 kb *scav-4* promoter or the promoter with the coding region of *scav-4* from the N2 genomic DNA and ligated them to the upstream of TagRFP and *unc-54* 3’UTR using Gibson Assembly. To create translational reporters for *scav-5* and *scav-6*, we cloned a 2 kb *scav-5* promoter with its coding region or a 2 kb *scav-6* promoter with its coding region to the upstream of TagRFP and an *unc-54* 3’UTR. To create the *ges-1p::cpt-4*, we ligated a 2 kb *ges-1* promoter to the coding region of *cpt-4* (cloned from N2 genomic DNA) and *unc-54* 3’UTR using Gibson Assembly. The constructs were then injected into the wild-type animals and several transgenic lines with extrachromosomal arrays were obtained and tested.

### CRISPR/Cas9-mediated gene editing

To recreate the L462F mutation in the *scav-4* gene in an otherwise wild-type background, we used 5’-CAGGCTAATGACTGAAAGAC-3’ as the Cas9 target sequence and single-stranded oligo 5’-CGGGAAGTACAATTGGTGGTGAATTTAGAATGATGTTT TCGATACCTGTGTTCCAATCGTTAGCCTGGACGTGAGTTAATTCCATAACTTAAAT-3’ as the repair template while introducing silence synonymous mutations in the target region to avoid cutting the repair template and facilitate genotyping. Recombinant spCas9 (Cat# M0646T, NEB) and single guide RNA synthesized using the sgRNA synthesis kit (Cat# E3322S, NEB) were injected together with the repair template into the wild-type animals. Transformants were screened for any successful editing using single-worm PCR. The correctly edited allele was named *unk59*.

To create knockout alleles of *scav-4*, we chose two Cas9 targets 5’-gatcaggATGAGGTTATTCG-3’ and 5’-ATCAGATGGTCAACTCCCAA-3’ (located in exon 1 and exon 2, respectively) and synthesized two guide RNAs using the NEB sgRNA synthesis kit. We injected them together with spCas9 and screened the F1 progeny for deletion at the beginning of the *scav-4* coding sequence and obtained *scav-4(unk49)* that deleted 146 nucleotides, including the entire intron 1 and parts of exon 1 and exon 2, and caused a frame shift. Similarly, to knock out *scav-6*, we chose two Cas9 targets 5’-GTTTGGAATTGTTACTTGGGCGG-3’ and 5’-CCGCTGCTCATTCAGGGgtaact-3’ (located in exon 1 and exon 5, respectively) and obtained the *scav-6(unk92)* allele that deleted 738 nucleotides (covering most of the first five exons) and caused frameshift.

### Bacterial feeding and lipid supplementation

In most experiments, *C. elegans* were fed with either *E. coli* OP50 or *Microbacterium sp.* JUb74. For lipid supplementation experiments, fatty acids were dissolved in ethanol and added to a 5 mL LB medium before inoculation. The bacteria culture was then allowed to grow overnight in an orbital incubator at 30°C (for JUb74) or 37°C (for OP50). Then the bacteria were concentrated to 500 µL before adding to the NGM plates.

For bacteria mixing experiments, the OD600 of the overnight cultures of OP50 and JUb74 were measured to estimate the amount of bacteria culture needed to achieve the desired ratios. After mixing the two bacteria cultures, the mixture is concentrated before being seeded onto NGM agar plates. After drying, the plates were irradiated by a UVP Crosslinker (CL-3000, Analytik Jena) for 20 minutes to kill the bacteria and prevent further growth.

### Fluorescent imaging and visualization of lipid droplets

Fluorescent imaging was performed on a Leica DMi8 inverted microscope with a Leica DFC7000 GT camera. To visualize lipid droplets in animals without LD markers, lipophilic fluorescent dye BODIPY™ 493/503, BODIPY™ 500/510 C1, C12, or BODIPY™ 558/568 C12 (Thermo Fisher Scientific) were diluted to a 1% solution in M9 buffer and then dropped on top of bacteria lawn before putting embryos onto the NGM plates. The animals were then transferred to 5 µl of M9 buffer on a 2% agarose pad with 3% 2,3-buantedione monoxime to sedate the animals. We imaged at least five day-one adult animals for each strain to measure the diameter of the lipid droplets visualized by the fluorescent dye using the Leica Application Suite X (LAS X) software. At least two biological replicates were done in each imaging experiment.

Fluorescence intensity is measured using the LAS X software to analyze the mean grey area encircling the entire body of the animals and normalized by subtracting the background intensity. For time-lapse imaging, animals were placed in 5 µl of 1:5 diluted 0.1 µm Polystyrene beads (Polysciences) on a 10% agarose pad to constrain the animals for a prolonged period. Images were taken once every 5 seconds for a 15-minute period to generate a time series.

### Lipid analysis

Total lipid profiles of worms were analysed using as previously reported protocol^61^ with some modifications. In brief, at least 10,000 adult worms were washed off from two to three 10 cm NGM plates with M9 buffer and placed into screw-capped centrifuge tubes to settle by gravity. The worm pellet was washed three times to remove all bacteria. Next, the worms were transferred to a glass tube by Pasteur pipette, with C17 fatty acid added as internal standard, and then incubated with 1 mL of 2.5% H2SO4 in methanol at 70°C for one hour to form fatty acid methyl esters (FAMEs). To terminate the reaction and extract the FAMEs, 1 mL of H2O and 1 mL of hexane were added to achieve phase separation after vortexing and centrifugation at 2000 g for 1 min. The top hexane layer was then transferred to a 2 mL screw top amber vial (Agilent). One mL of hexane was added again to maximize the FAME extraction from the glass tube. Hexane in the gas chromatography-mass spectrometry (GC-MS) compatible vials were then dried under a stream of N2 gas. To resuspend the FAMEs, 50 µL of hexane was added and transferred into a vial insert (Agilent). One µL of the sample was analyzed by GC with an Agilent 7890B GC system equipped with a 100 m × 0.25 mm SP-2560 column (Supelco), helium as the carrier gas at 1.2 mL/min, and a flame ionization detector. The GC was programmed for an initial temperature of 100°C for 5 min followed by an increase of 4°C/min to 240°C. Scan mode of mass spectra from m/z 50 to 550 was used to scan all ions in the mass spectrometer (Agilent 7010 Triple Quadrupole Mass Spectrometer). Qualitative analysis of the FAMEs are confirmed by comparing with 37 Component FAME Mix (Supelco) or Bacterial Acid Methyl Ester (BAME) Mix (Supelco).

To analyze fatty acids from bacteria, 5 mL of overnight culture of bacteria at 30°C (for JUb74) or 37°C (for OP50) in LB media was pelleted by centrifuging at 3000 g for 10 minutes. The bacteria pellets were then washed with water at least three times before transferring them to a clean glass for transmethylation.

To extract total lipid for thin layer chromatography, at least 10,000 worms were collected and were lysed in 500 µL of RIPA buffer [25 mM Tris-HCl, 100 mM NaCl, 1 mM EDTA, 0.5% NP-40, 1 mM PMSF, 1 mM Na3VO4, 1mM NaF, and protease inhibitor cocktail (Roche)] before being homogenized by sonication (Branson Sonifier 250). The lysate was then centrifuged at 5000 g for 10 minutes at 4°C. 400 µL of the supernatant was added to ice-cold Folch solution (2:1 v/v chloroform: methanol) for lipid extraction, and 100 µL was used to determine protein concentration using BCA Protein Assay Kit (Thermo Fisher). After one hour of shaking in the ice-cold Folch solution, the mixture was centrifuged at 5000 rpm for 5 minutes and the bottom chloroform layer was transferred by a Pasteur pipette to a clean GC glass vial. The collected chloroform was then dried under a stream of N2 gas and resuspended in 50 µL chloroform. The extracted lipid and triglyceride standard (Merck) were then added to a silica gel HL uniplates (Analtech) and developed by a mixture of organic solvents (hexane: dimethyl ether: acetic acid 80:20:3, v/v). Separated lipids were visualized by iodine vapor inside a closed chamber with silica gel and iodine crystals and then scraped off for transmethylation.

Fatty acid quantifications were performed at the Centre of PanorOmic Sciences at the University of Hong Kong according to a previous protocol^62^. 100 μl of chloroform with 20 μg C19:0 fatty acid internal standard was added to the sample. Worm samples were extracted with 5 rounds of 2:1 chloroform/methanol, followed by sonication. After centrifugation, the supernatant was further cleaned by liquid-liquid extraction in 0.73% NaCl and methanol. The resultant mixture was dried under a stream of N2 gas at 45°C before transesterification and thereafter. 1 ml of methanol and 50 μl of concentrated hydrochloric acid (35%, w/w) were added to the sample. The solution was overlaid with N2 and the tube was tightly closed. After vortexing, the tube was heated at 100°C for 1.5 h. Once cooled to room temperature, 1 ml of hexane and 1 ml of water were added for FAMEs extraction. The tube was vortexed and after phase separation, 1 μl the hexane phase was injected for GC-MS analysis using an Agilent 7890B GC - Agilent 7010 Triple Quadrupole Mass Spectrometer system in SCAN and SIM mode. The sample was separated through an Agilent DB-23 capillary column (60 m × 0.25 mm ID, 0.15 μm film thickness) under constant pressure of helium at 33.4 psi. The GC oven program started at 50°C (hold time 1 min) and was increased to 175°C at a ramp rate of 25°C/min. The temperature was then raised to 190°C (hold time 5 min) at a ramp rate of 3.5°C/min. Finally, the temperature was raised to 220°C (hold time 4 min) at a ramp rate of 2°C/min. Inlet temperature and transfer line temperature were 250°C and 280°C, respectively. Characteristic fragment ions were monitored in SIM mode throughout the run. Mass spectra from m/z 50-350 were acquired in SCAN mode. Data analysis was performed using the Agilent MassHunter Workstation Quantitative Analysis Software. Linear calibration curves for each analyte were generated by plotting the peak area ratio of external/internal standard against standard concentration at different concentration levels. Analytes were confirmed by comparing the ratio of characteristic fragment ions in the sample and standard.

### Transcriptomic analysis

Total RNA from N2, CGZ413 *scav-4(unk59; L462F)*, and CGZ337 *scav-4(unk49)* fed with OP50 or JUb74 were extracted using TRIzol reagent (Thermo Fisher). Samples were sent to BGI (Beijing Genome Institutes, Shenzhen, China) for standard library construction and pair-end sequencing. Around 20 million reads were obtained for each sample, and the reads were aligned to the *C. elegans* genome (WS235) using STAR 2.7. To identify genes differentially expressed, the transcript counts were analyzed using DESeq2, and genes with false discovery rate–adjusted *p* values below 0.05 and log2(fold change) above 2 or below -2 were identified. Gene enrichment analysis was then performed on the genes showing significant changes in expression levels using the DAVID functional annotation tool (2021 version). The raw RNA-seq data were uploaded to NCBI Gene Expression Omnibus (GEO) with the accession number GSE282709.

### Lifespan, growth rate, and brood size determinations

The lifespans of the animals were determined by recording the survival of 20 synchronized L4 animals on an NGM plate with OP50 or JUb74 every day until all 20 original worms were dead. For the first five days of the experiments, the worms were transferred to a fresh NGM plate with the same bacteria to separate the testing subjects from their progenies. Animals that died from abnormal reasons, such as dehydration due to moving away from the NGM agar to the Petri dish wall were not counted in the final percentage. Three replicates were performed, and mean percentages were shown. All data were combined for the calculation of the statistical significance of the difference between groups using the log-rank (Mantel-Cox) method.

Growth rates of the animals were measured by observing the developmental stages of at least 50 animals in each group of animals or diet at 72 hours and 96 hours after placing the bleach-synchronized embryos onto the NGM plates with bacteria. At least three replicates were performed in each group.

Brood sizes of the animals were calculated by the total amount of embryos laid by the same hermaphrodite until it reached the end of its reproductive period. At least ten animals were analyzed in each group. Each animal was grown in individual NGM plates with bacteria and transferred to a new NGM plate every day; the number of embryos on the plate was counted after the transfer. Larvae from the old plates were checked two days later to determine the number of hatched animals.

### ATP production measurement

Worms fed with different bacteria diets are harvested and homogenized by sonication. ATP concentrations were measured using the ATP Determination Kit (Thermo Fisher Scientific). ATP concentrations were then normalized to protein concentrations measured by BCA Protein Assay Kits (Thermo Fisher Scientific). Both bioluminescence and colorimetric measurements were done on SpectraMax iD3 and iD5 Multi-Mode Microplate Readers (Molecular Devices).

### Phenotypic scoring and statistical analysis

All data were plotted as mean ± SD using GraphPad Prism 8.0. For multiple comparisons, we performed post-ANOVA Dunnett’s multiple comparisons test of different treatment groups with a single control group or a Tukey’s HSD test for all pairs of groups. Student’s *t-*test was used to compare only two groups. Single asterisks indicate statistical significance at *p* < 0.05, and double asterisk for *p* < 0.01.

## Supporting information

Supplemental Figures S1-S7 and legends for other supplemental materials

## ACKNOWLEDGEMENT

We thank Prof. Ho Yi Mak at the Hong Kong University of Science and Technology and Prof. Aixin Yan at the University of Hong Kong for sharing reagents and equipment and helpful discussion. We thank Terence Lee in the Zheng lab for technical assistance. The graphical abstract was created using BioRender.com. This study is supported by funds from the National Natural Science Foundation of China (Excellent Young Scientists Fund for Hong Kong and Macau, 32122002), the Health Bureau of Hong Kong (HMRF 09201426 to C.Z.), and the Research Grant Council of Hong Kong (GRF 17105523, GRF 17106322, GRF 17107021, and CRF C7026-20G). Some strains used in this study were provided by the Caenorhabditis Genetics Center, which is funded by the NIH Office of Research Infrastructure Programs (P40 OD010440).

## AUTHOR CONTRIBUTIONS

C.Z. and K.U.R.L. conceived the study and wrote the manuscript. K.U.R.L., W.K.L., and M.Y. carried out most of the experiments. C.W. did some initial characterization of the phenotype. L.C.Y. performed the RNA-seq analysis, J.C.-Y.L. provided resources and technical support. C.Z. secured funding and supervised the project. All authors read and approved the manuscript.

