## Supplemental Figures S1-S7 and legends for other supplemental materials for "Branched chain fatty acid-rich diet promotes lipid droplet enlargement and impacts organismal health in *Caenorhabditis elegans*"

**Supplemental Information**
**Figure S1. JUb74 diet induces enlarged LDs.**
**Figure S2. Fatty acid profiles of OP50 and JUb74 mixture and lipid-supplemented bacteria.**
**Figure S3. JUb74 diet-induced LDs are resistant to hydrolysis and prone to fusion.**
**Figure S4. L462F mutation in *scav-4* promotes the formation of supersized LDs under the JUb74 diet.**
**Figure S5. Loss of *scav-4* reduces the absorption of dietary lipids.**
**Figure S6. Mutations in *scav-4* cause transcriptomic changes under the JUb74 diet.**
**Figure S7. JUb74-induced upregulation of *fat-7* requires both metabolic regulator NHR-49/NHR-80 and mitoUPR activator ATFS-1.**

**Movie S1. Fusion of two large LDs in animals fed with JUb74.** LIU1 *ldrIs1[dhs-3p::dhs-3::GFP + unc-76(+)]* animals fed with JUb74 were imaged for 5 minutes. The video is played at 15x speed.
**Movie S2. Fusion between one large and one small LD in animals fed with JUb74.** LIU1 *ldrIs1[dhs-3p::dhs-3::GFP + unc-76(+)]* animals fed with JUb74 were imaged for 5 minutes. The video is played at 15x speed.
**Movie S3. Frequent LD fusion in *scav-4(L462F)* mutants fed with JUb74.** LDs were visualized using green BODIPY. Multiple fusion events were observed in the 5-minute clip, which is played at 15x speed. In the video, five LDs fused into one big one within 2 minutes.

**Table S1. Genes differentially expressed between wild-type and *scav-4(unk59; L462F)* animals fed with OP50.**
**Table S2. Genes differentially expressed between wild-type and *scav-4(unk59; L462F)* animals fed with JUb74.**
**Table S3. Genes differentially expressed between wild-type and *scav-4(unk49; deletion)* animals fed with OP50.**
**Table S4. Genes differentially expressed between wild-type and *scav-4(unk49; deletion)* animals fed with JUb74.**
**Table S5. Genes differentially expressed between wild-type animals fed with OP50 and JUb74.**
**Table S6. Genes differentially expressed between *scav-4(unk59; L462F)* animals fed with OP50 and JUb74.**
**Table S7. Strains, DNA, and primers used in this study.**

**Supplemental File 1. Annotated JUb74 genome in the genebank format.**
**Supplemental File 2. JUb74 genome sequence in the fasta format.**
**Supplemental File 3. The 16S rRNA sequence of JUb76.**
**Supplemental File 4. The 16S rRNA sequence of JUb84.**
**Supplemental File 5. The 16S rRNA sequence of JUb90.**

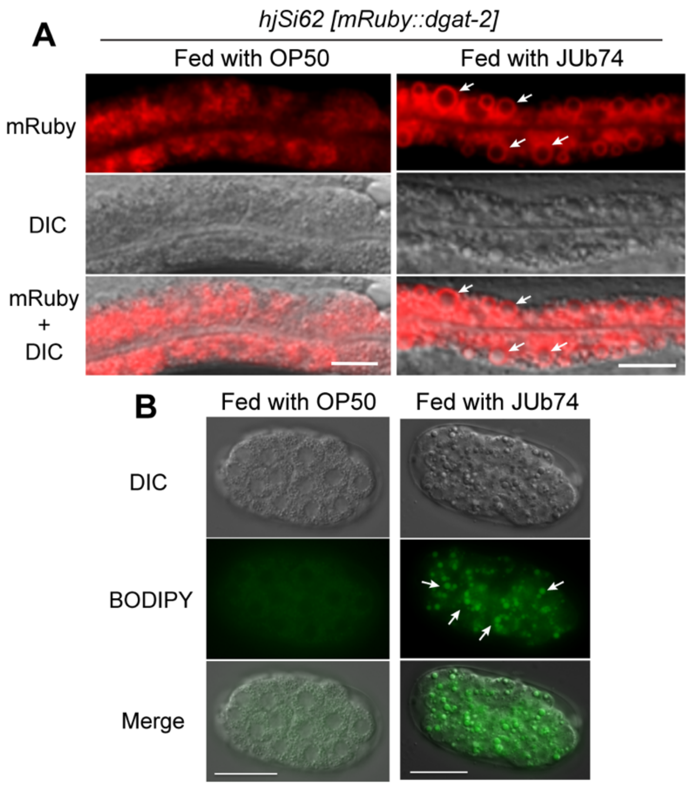


**Figure S1. JUb74 diet induces enlarged LDs.** (A) Animals carrying the *mRuby::dgat-2* transgene were fed with either OP50 or JUb74, and the surface of the enlarged LDs was labeled by the red fluorescent signal (arrows). (B) Eggs (~26-cell stage) laid by animals fed with either OP50 or JUb74 and treated with BODIPY dye. Arrows point to the large LDs in the embryos. Scale bar = 20 µm. The eggs extracted from animals fed with OP50 were imaged using 500-ms exposure time, while eggs from animals fed with JUb74 were imaged using 182-ms exposure time because the BODIPY signal was too strong to show the individual LDs.

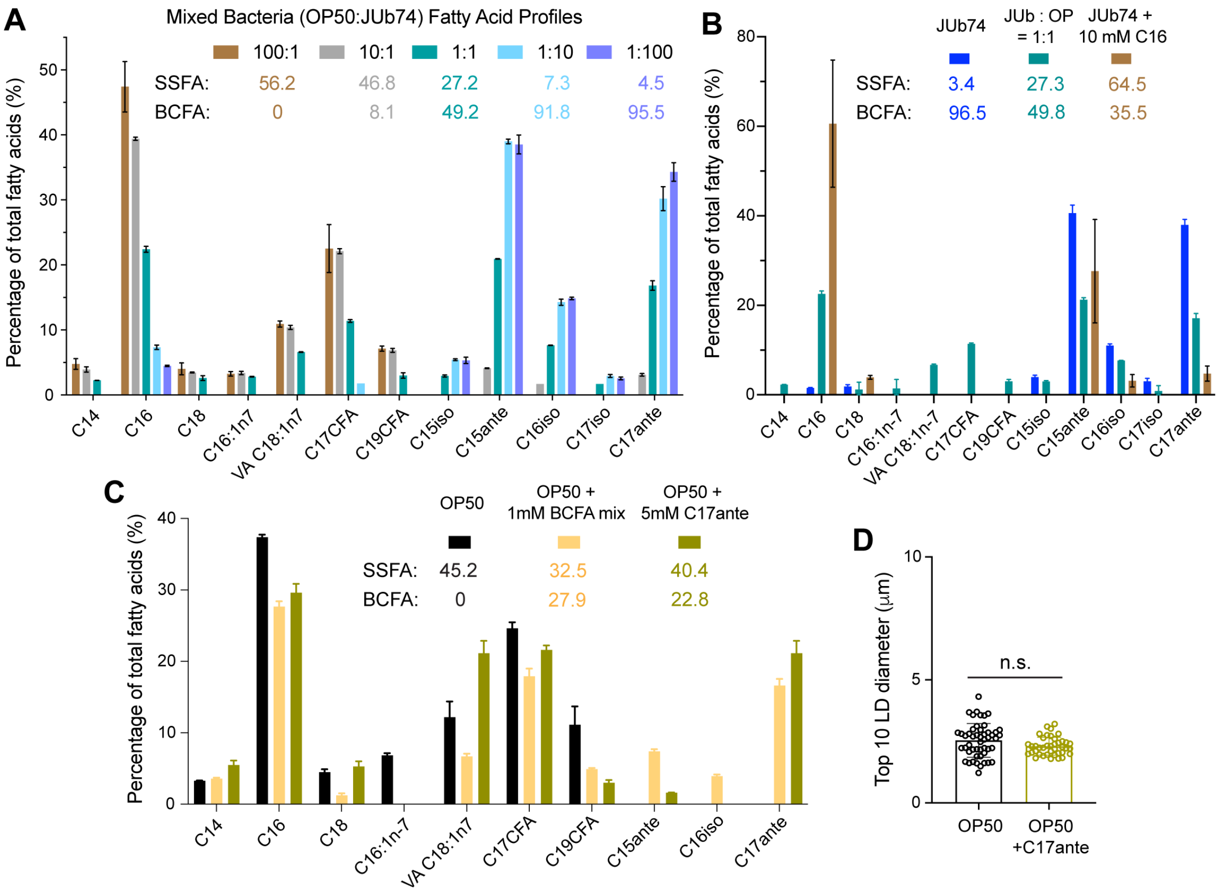


**Figure S2. Fatty acid profiles of OP50 and JUb74 mixture and lipid-supplemented bacteria.** (A) Fatty acid profiles of bacterial mixture between OP50 and JUb74 at different ratios. Combined percentages of SSFAs and BCFAs were shown. (B) Fatty acid profiles of JUb74, a one-to-one mixture of JUb74 and OP50, and JUb74 supplemented with 10 mM palmitic acid (C16). (C) Fatty acid profiles of OP50 supplemented with a mixture of in total 1 mM BCFAs (0.4 mM C15anteiso, 0.4 mM C17anteiso, and 0.2 mM C16iso) or 5 mM C17 anteiso. (D) The diameter of the ten biggest LDs in animals fed with OP50 or OP50 supplemented with 5mM C17anteiso; “n.s.” means statistically not significant in a *t*-test.

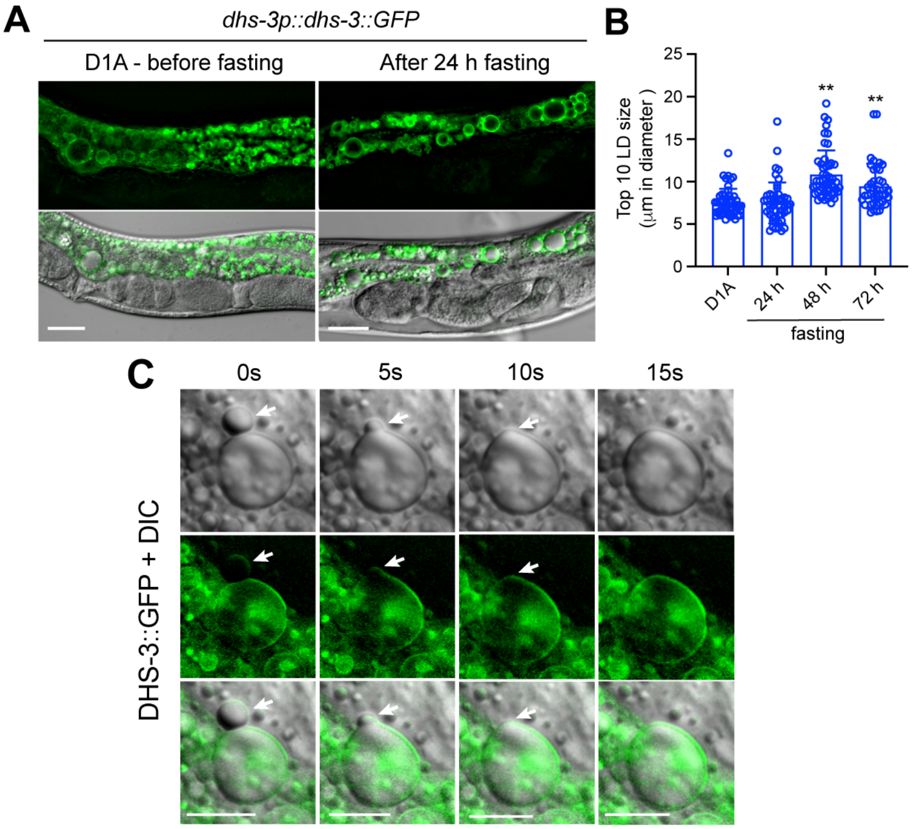


**Figure S3. JUb74 diet-induced LDs are resistant to hydrolysis and prone to fusion.** (A) LD labeled by the marker *dhs-3p::dhs-3::GFP* in animals fed with JUb74 from egg to the day-one adult (D1A) stage and then starved for 24, 48, and 72 hours. Scale bar = 20 µm. (B) The diameter of the ten biggest LDs in animals before and after fasting. Double asterisks indicate $p < 0.05$ in a post-ANOVA Dunnett's test in comparison with the day-one adults before fasting. The increased LD size is likely a result of continuous fusion of preexisting LDs before the fasting. (C) Time-lapse images that track the fusion of a big LD with a smaller one (indicated by the arrows) in the LIU1 *ldrIs1[dhs-3p::dhs-3::GFP]* strain that labeled the LD surface. Scale bar = 10 µm.

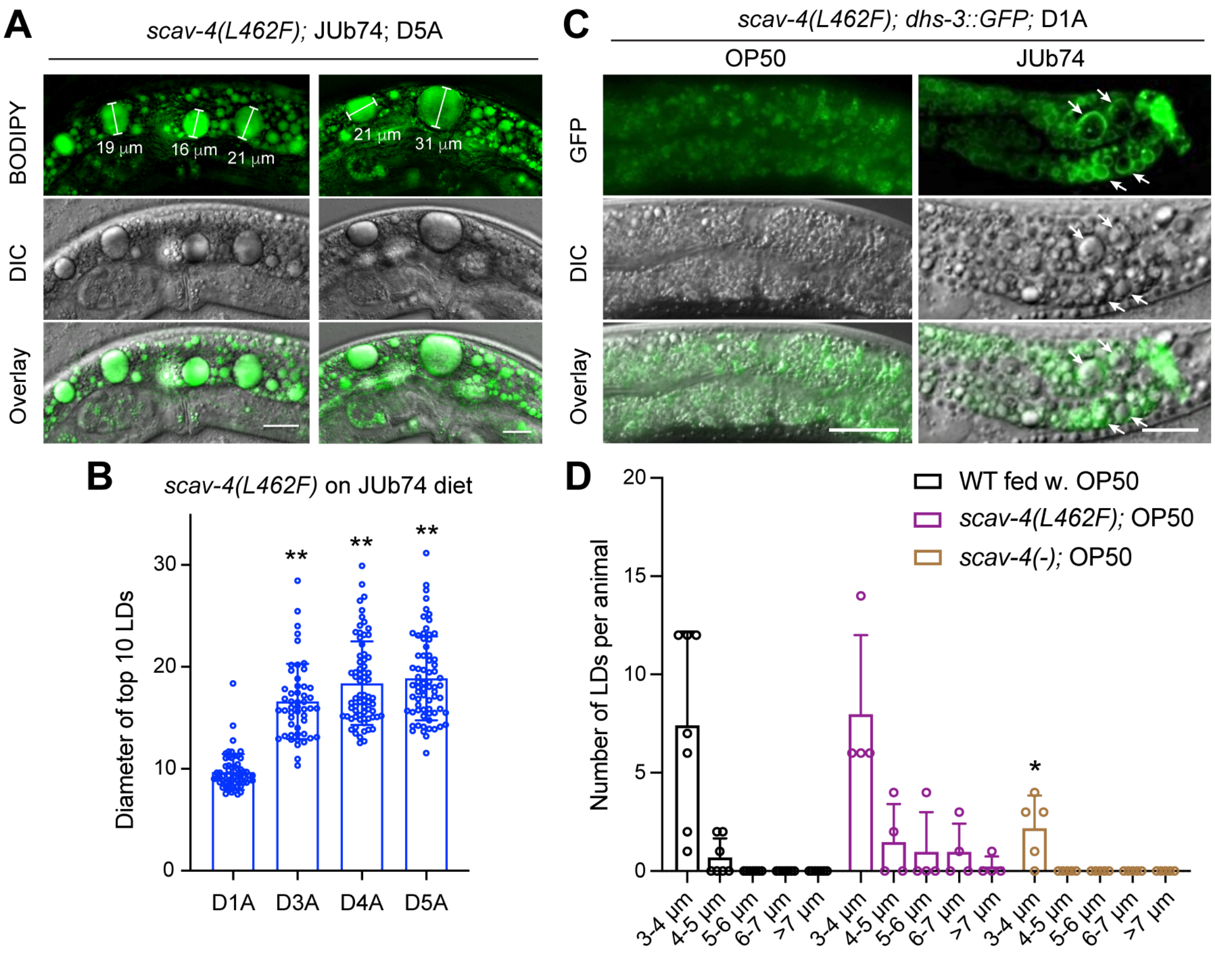


**Figure S4. L462F mutation in *scav-4* promotes the formation of supersized LDs under the JUb74 diet.** (A) *scav-4(L462F)* mutants were fed with the JUb74 diet and supplemented with BODIPY at day-five adult stage. Supersized LDs with diameters larger than 15 µm are labeled; scale bar = 20 µm. (B) The diameter of the ten biggest LDs in *scav-4(L462F)* mutants fed with JUb74 at different adult stages. Double asterisks indicate $p < 0.01$ in a post-ANOVA Dunnett's test. (C) *scav-4(L462F)* mutants carrying the LD marker *ldrIs1[dhs-3p::dhs-3::GFP]* were fed with JUb74 and imaged at day-one adult stage. Arrows indicate enlarged LDs. (D) The distribution of LD diameters in wild-type, *scav-4(unk59; L462F)*, and *scav-4(unk49)* animals fed with OP50 at the day-one adult stage. A single asterisk indicates $p < 0.05$ in a *t*-test in comparison with the wild-type animals for the same size category.

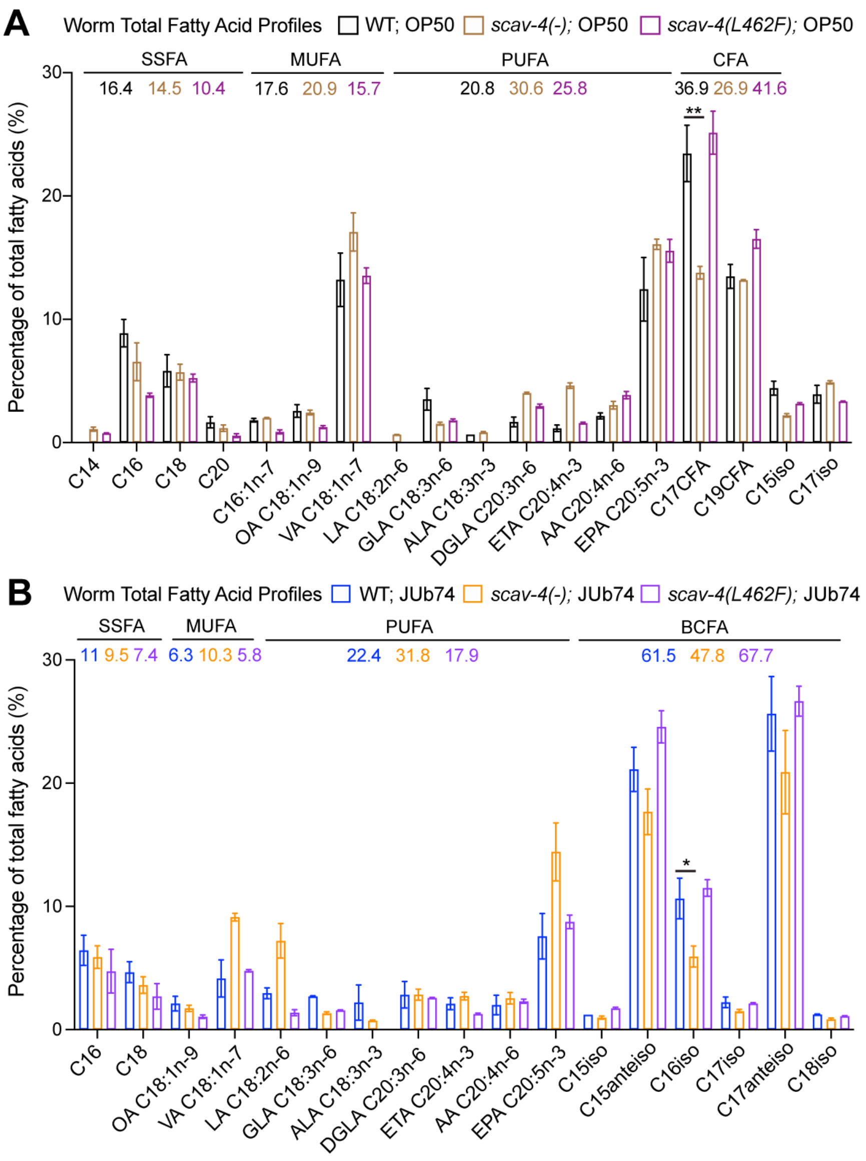


**Figure S5. Loss of *scav-4* reduces the absorption of dietary lipids.** (A) Fatty acid profiles of wild-type, *scav-4(unk49)*, and *scav-4(unk59; L462F)* animals fed with OP50. The combined percentages of SSFAs, MUFAs, PUFAs, and CFAs were shown. Double asterisks indicate $p < 0.01$ in a *t*-test. (B) Fatty acid profiles of wild-type, *scav-4(unk49)*, and *scav-4(unk59; L462F)* animals fed with JUb74. The combined percentages of SSFAs, MUFAs, PUFAs, and BCFAs were shown. Single asterisk indicates $p < 0.05$ in a *t*-test.

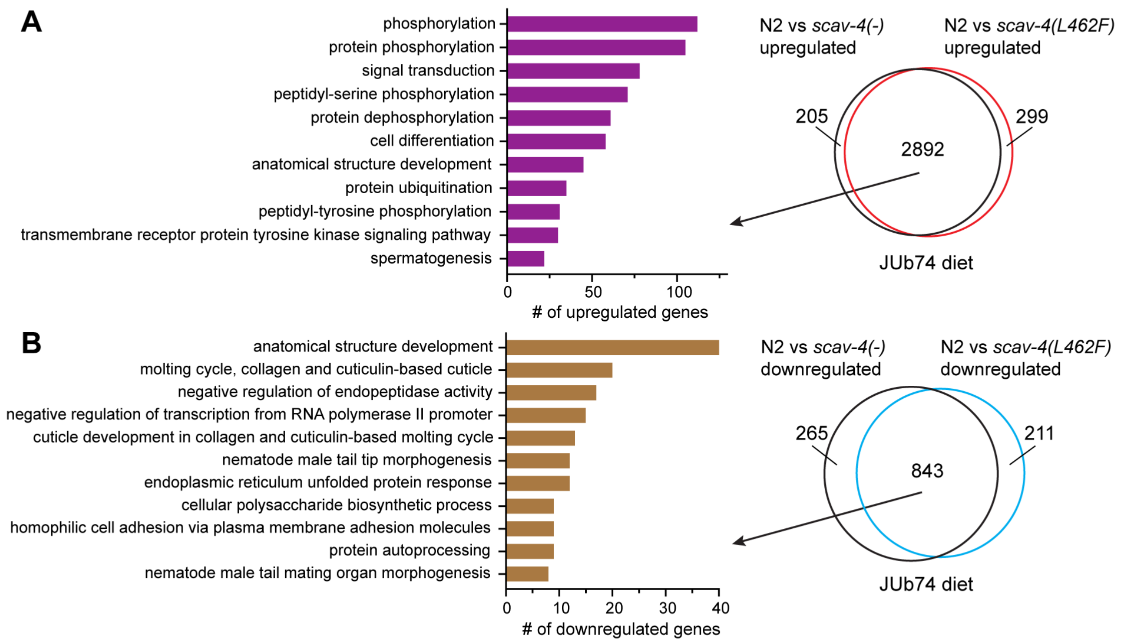


**Figure S6. Mutations in *scav-4* cause transcriptomic changes under the JUb74 diet.** (A) Genes upregulated in *scav-4(unk49)* and *scav-4(unk59; L462F)* animals compared to the wild-type animals when fed with JUb74. The 2,892 overlapping genes were subjected to gene ontology (GO) analysis. Significantly enriched GO terms (adjusted $p < 0.05$) were listed together with the number of upregulated genes associated with the term. (B) Genes downregulated in *scav-4(unk49)* and *scav-4(unk59; L462F)* animals compared to the wild-type animals when fed with JUb74. The 843 overlapping genes were subjected to analysis. Significantly enriched GO terms (adjusted $p < 0.05$) were listed together with the number of downregulated genes associated with the term.

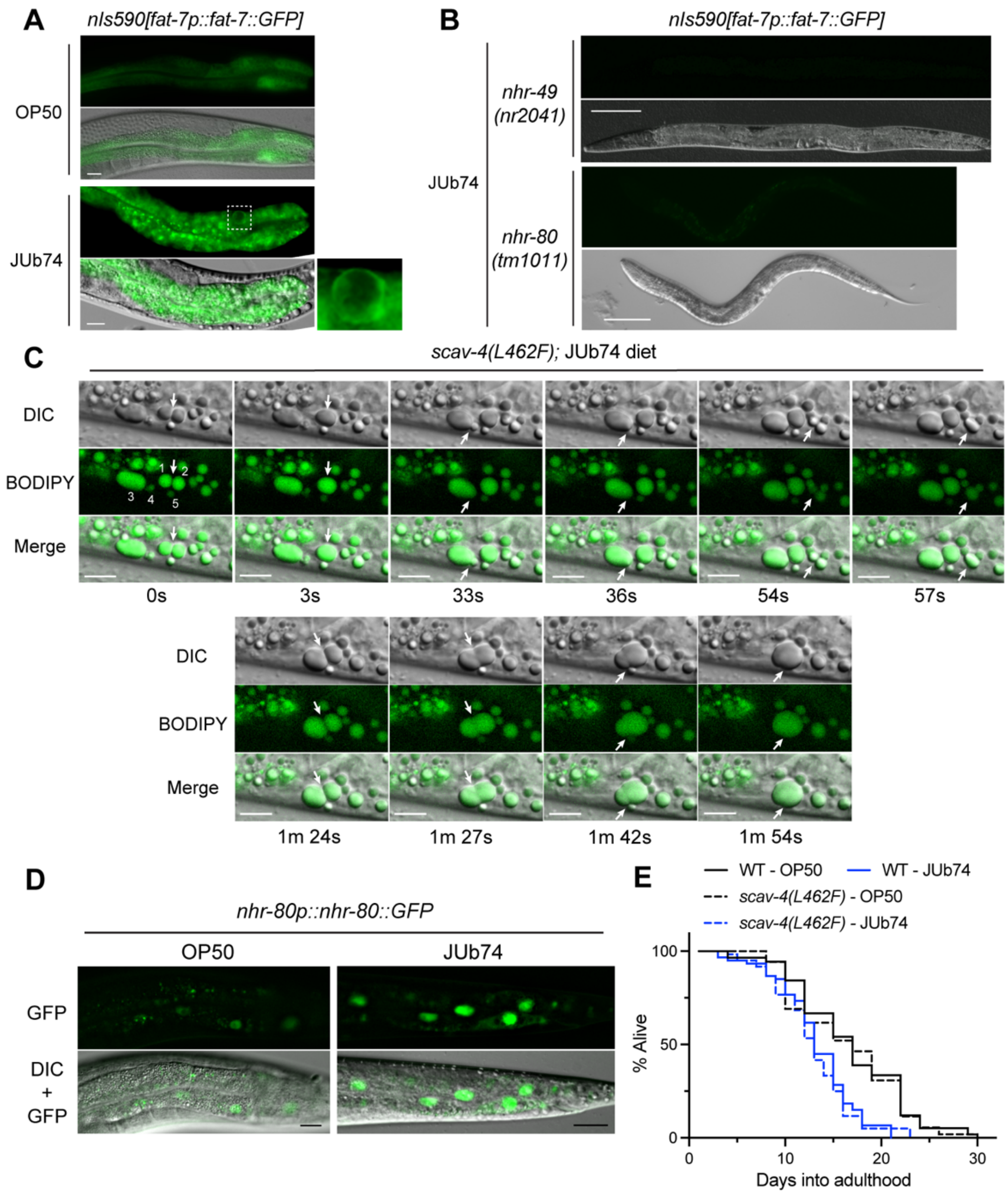


**Figure S7. JUb74-induced upregulation of *fat-7* requires both metabolic regulator NHR-49/NHR-80 and mitoUPR activator ATFS-1.** (A) Representative images of DMS303 *nIs590[fat-7p::fat-7::GFP]* animals fed with OP50 and JUb74. The dashed box is enlarged to show that FAT-7::GFP is enriched on the surface of LDs under the JUb74 diet. Scale bar = 20 µm. (B) The loss of *fat-7p::fat-7::GFP* expression in *nhr-49(-)* and *nhr-80(-)* mutants under JUb74 diet. Scale bar = 100 µm. (C) LD fusion in *scav-4(L462F)* mutants fed with JUb74. Selective frames were shown in a time-lapse series. Arrows indicate fusion events. The numbers 1~5 in the first GFP image label the five LDs that fused together to form one large LD (arrow in the last frame). Scale bar = 10 µm. (D) The expression of *lynEx1[nhr-80p::nhr-80::GFP]* under OP50 and JUb74 diet. (E) Survival curves of wild-type and *scav-4(unk59; L462F)* animals under OP50 and JUb74 diets.
